# On growth and form of the mammary gland: Mesenchyme instructs growth while epithelium directs branching

**DOI:** 10.1101/2023.04.24.538064

**Authors:** Qiang Lan, Ewelina Trela, Riitta Lindström, Jyoti Satta, Beata Kaczyńska, Mona M. Christensen, Martin Holzenberger, Jukka Jernvall, Marja L. Mikkola

## Abstract

Mammary gland is a unique organ that undergoes dynamic alterations throughout a female’s reproductive life, making it an ideal model for developmental, stem cell and cancer biology research. Mammary gland development begins *in utero* and proceeds via a quiescent bud stage before the initial outgrowth and subsequent branching morphogenesis. How mammary epithelial cells transit from quiescence to an actively proliferating and branching tissue during embryogenesis and, importantly, how the branch pattern is determined remain largely unknown. Here we provide evidence indicating that epithelial cell proliferation, segregation into basal and luminal lineages that characterize the postnatal mammary duct, and onset of branching are independent processes, yet partially coordinated by the Eda signaling pathway. By performing heterotypic and heterochronic epithelial-mesenchymal recombination experiments between mammary and salivary gland tissues and *ex vivo* live imaging, we demonstrate that unlike previously concluded, the mode of branching is an intrinsic property of the mammary epithelium while the growth pace and density of the mammary ductal tree are governed by the mesenchyme. Transcriptomic profiling and *ex vivo* and *in vivo* functional studies disclose that mesenchymal Wnt/ß-catenin signaling, and in particular IGF-1 downstream of it critically regulate mammary gland growth. These results underscore the general need to carefully decompose the different developmental processes producing branched organs.

## Introduction

Branching morphogenesis is a common developmental process driving the formation of a number of organs including lung, kidney, salivary and mammary gland ^1^. Although some fundamental principles are shared, each organ employs its unique branching strategy – mode and density of branching – to achieve the proper architecture tailored to its function ^1–3^. In recent decades, significant advancements have been made in unraveling the underlying mechanisms of branching morphogenesis in various organs and species. However, many questions remain unanswered, especially regarding the mammary gland as much of the research focus has been on its postnatal growth^1, 2^. Yet, mammary gland morphogenesis commences already during fetal life by formation of placodes, local epithelial thickenings, in the flanks of the fetus. How these early steps of branching morphogenesis differ between mammary gland and other organs remains poorly understood.

In mice, five pairs of mammary placodes emerge around embryonic day 11 (E11). Placodes invaginate by E13 giving rise to buds that are now surrounded by condensed, mammary-specific mesenchyme ^4–6^. Mammary buds stay relatively non-proliferative until E15-E16 when they sprout toward the adjacent ‘secondary’ mammary mesenchyme, the fat pad precursor tissue that later gives rise to the adult stroma. Branching begins at E16, and by E18 (1-2 days prior to birth) mammary rudiments have developed into small ductal trees with 10-25 branches ^3, 7^. In contrast to the postnatal bilayered mammary epithelium consisting of outer basal and inner luminal cells, embryonic mammary rudiments undergo branching as a solid mass of epithelial cells without lumen. Mammary rudiments initially consist of multipotent precursors that become restricted to basal and luminal lineages during later stages of embryogenesis ^8, 9^. The mechanisms governing the exit from quiescence and acquisition of branching ability are still enigmatic. The observation that the initial outgrowth coincides with activation of proliferation and lineage segregation has led to the hypothesis that they might be functionally connected ^8, 10^. However, whether a causal link exists between these phenomena is currently unknown.

Reciprocal epithelial-mesenchymal tissue interactions are critical for mammary gland development at all stages. Many signaling pathways essential for mammary placode and bud formation have been identified, but the paracrine factors regulating branching during embryogenesis are less well understood ^4, 5, 11, 12^. The tumor necrosis factor family member ectodysplasin A1 (Eda) is one such mesenchymal factor: Eda deficiency compromises ductal growth and branching, while mice overexpressing Eda exhibit a dramatic ductal phenotype with precocious sprouting and excessive growth and branching^13, 14^. In addition, the Wnt and fibroblast growth factor (Fgf) pathways are likely involved ^7, 11^, but the early developmental arrest observed in mice where these pathways are inactivated ^15, 16^ has hampered elucidation of their exact roles during branching morphogenesis.

Importantly, the current paradigm posits that the mesenchyme specifies the epithelial branching pattern in all branched organs ^1, 3^. This conclusion stems from tissue recombination experiments where epithelia and mesenchymes of different origins have been exchanged: lung mesenchyme instructs the kidney epithelium to adopt a lung-type branching pattern while organ-specific mode of branching is maintained in homotypic tissue recombinants ^17, 18^. The same conclusion was drawn from the pioneering experiments involving salivary gland mesenchyme and mammary gland epithelium. Even though the mammary epithelium retained its cellular identity, the branch pattern was reported to be salivary gland-like: branches formed at higher density and by tip clefting rather than lateral branching ^19, 20^. In addition, salivary gland mesenchyme promoted much faster growth. Although the evidence from the early experiments appear compelling, the underlying molecular basis remained elusive.

To uncover the regulation of mammary gland branching, we first revisited the heterochronic tissue recombination using mammary tissues. Our results show that the timing of the initial branching is epithelium-dependent, yet epithelial-mesenchymal interactions are indispensable for the outgrowth to occur. Further analysis suggests that onset of branching is independent of onset of proliferation and lineage segregation, yet they are partially coordinated by the Eda pathway. In strong contrast to the previous reports and to the paradigm of the role of the mesenchyme in directing branching ^19, 20^, live imaging disclosed that salivary gland mesenchyme failed to switch the mode of mammary branching into salivary-like. This implies that branch pattern formation is an intrinsic property of the mammary epithelium. Nevertheless, salivary mesenchyme had a major growth-promoting effect on the mammary epithelium once it had acquired branching capacity. Transcriptomic profiling of mammary and salivary gland mesenchymes identified mesenchymal Wnt/ß-catenin pathway and its downstream target *Igf1* as potential drivers of epithelial growth, thereby decomposing mode of branching from growth control in mammary development.

## Results

### The timing of onset of branching is an intrinsic property of the mammary epithelium

To assess whether timing of the mammary initial branching can be influenced by tissues of different developmental stages, we performed heterochronic epithelial-mesenchymal recombination experiments. To this end, we used tissues micro-dissected from fluorescently labeled transgenic mice allowing day-to-day imaging, as well as evaluation of the purity of the tissue compartments (Fig. 1a). Because anterior mammary glands are more advanced in their development than the posterior ones ^7^, only mammary glands 1 to 3 were used throughout the study, unless otherwise specified, to avoid any biases caused by the asynchrony.

**Fig. 1.**
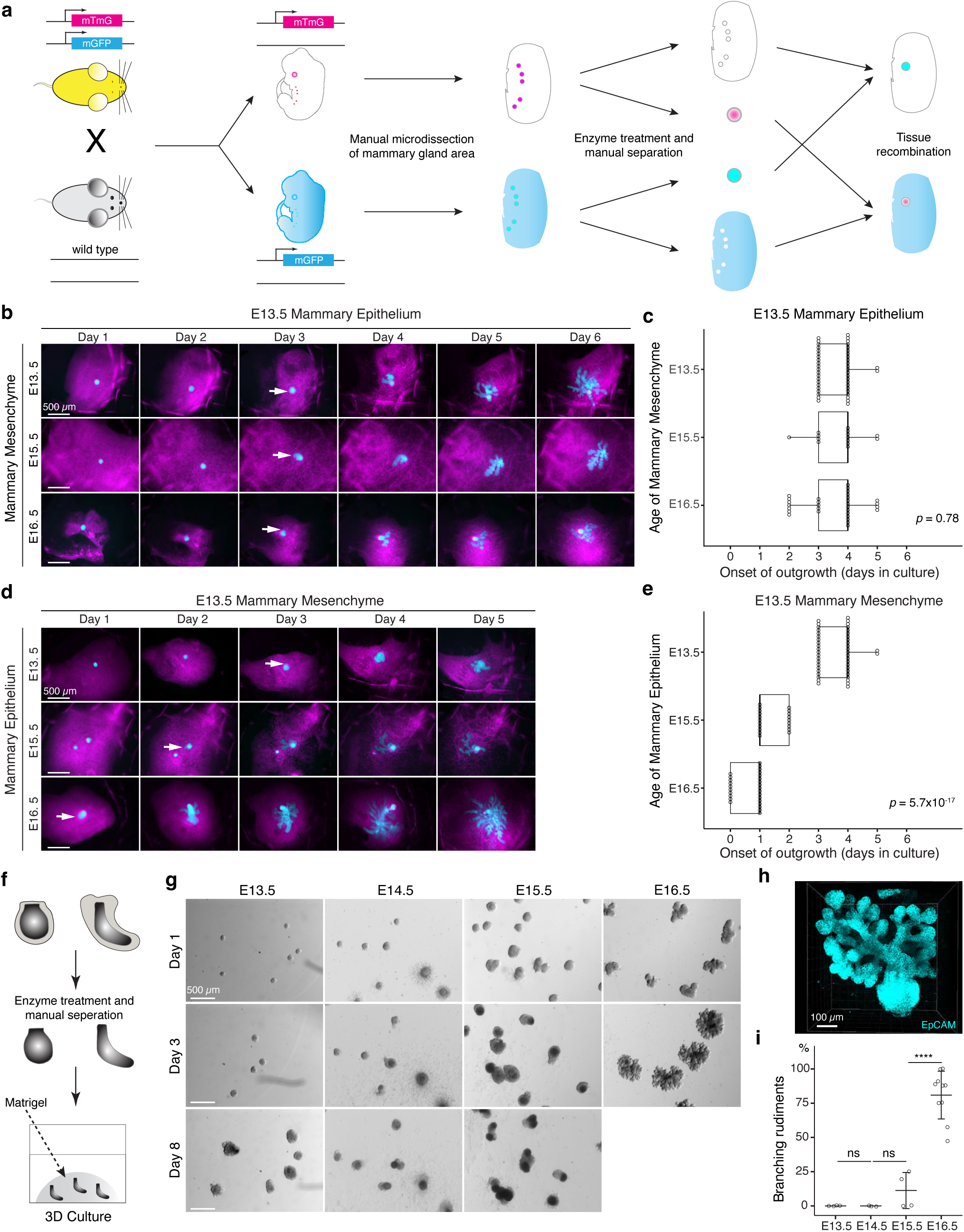
The timing of mammary gland outgrowth is an inherent property of the epithelium. **a**, A scheme illustrating the experimental procedure used in tissue recombination experiments. **b**, Representative images showing the onset of outgrowth of E13.5 mammary epithelia recombined with E13.5, E15.5, or E16.5 mammary mesenchymes, respectively. The appearance of the primary outgrowth is indicated with arrow. Scale bar, 500 µm. **c**, Quantification of the time (in days) required for onset of the branching. Data were pooled from 3-6 independent experiments of E13.5 mammary epithelia recombined with E13.5 (n=46 explants), E15.5 (n=14), and E16.5 (n=30) mammary mesenchymes. Statistical significance was assessed with the Kruskal–Wallis test. **d**, Representative images showing onset of outgrowth of E13.5, E15.5 and E16.5 mammary epithelia recombined with E13.5 mammary mesenchymes. The appearance of the primary outgrowth is indicated with arrow. Scale bar, 500 µm. **e**, Quantification of the time (in days) required for the onset of the primary outgrowth. Data were pooled from 3-6 independent experiments of E13.5 (n=46 explants), E15.5 (n=20) and E16.5 (n=27) mammary epithelia recombined with E13.5 mammary mesenchyme. Statistical significance was assessed with the Kruskal–Wallis test. **f**, A scheme illustrating the 3D culture of intact, mesenchyme-free epithelial mammary rudiments. **g**, Representative images showing the growth of E13.5, E14.5, E15.5 and E16.5 epithelial mammary rudiments in 3D culture; only E16.5 mammary rudiments were capable of branching (see also Supplementary Fig. 1). Scale bar, 500 µm. **h**, Representative 3D projection image of an EpCAM-stained E16.5 mammary rudiment after three days of 3D culture in Matrigel. Scale bar, 100 µm. **i**, Quantification of branching mammary rudiments in 3D culture. Data are presented as percentage of branching mammary rudiments (mean ± SD) from a total of 4 (E13.5), 3 (E14.5), 4 (E15.5), and 10 (E16.5) independent experiments (each with minimum 6 rudiments in culture). The statistical significances were assessed using unpaired two-tailed Student’s *t*-test with Bonferroni correction. ns, non-significant; ****, *p* < 0.001.

It has been previously shown that early (E12) mammary mesenchyme does not alter the onset of branching of the mammary epithelium (E12 to E16) in *ex vivo* tissue recombination experiments ^19^. However, the ability of late mammary mesenchyme to advance epithelial outgrowth and branching has not been assessed. To answer this question, we recombined E13.5 mammary epithelia (quiescent bud stage) with E13.5, E15.5, or E16.5 (when the very first branches are evident) mammary mesenchymes. In the control explants (E13.5 epithelia with E13.5 mesenchyme), branching started after 3-4 days of culture (Fig. 1b,c), in good agreement with development *in vivo*. No precocious branching was observed when ‘older’ mesenchyme was used: when E13.5 epithelia were cultured with either E15.5 or E16.5 mesenchyme, branching was again evident only after 3-4 days of culture (Fig. 1b, c). As an additional control, we performed similar experiments as described by Kratochwil ^19^, and cultured E13.5, E15.5 or E16.5 mammary epithelia with E13.5 mammary mesenchyme (Fig. 1d). As previously reported, all epithelia branched in E13.5 mesenchyme, and outgrowth started after 3-4, 1-2, and 0-1 days of culture, respectively (Fig. 2e), correlating with the stage of epithelium and its developmental pace *in vivo*.

**Fig.2.**
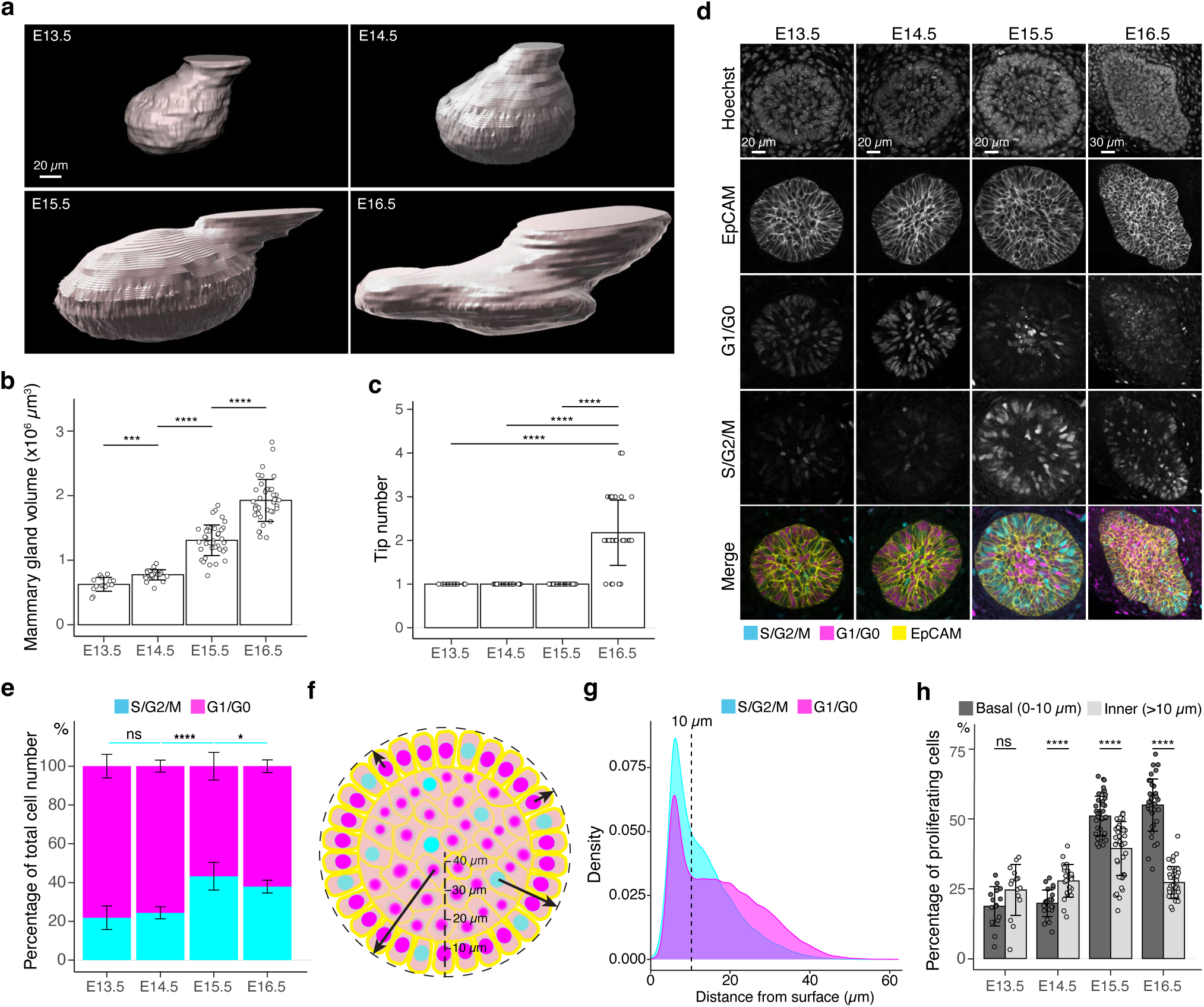
Cell cycle dynamics in embryonic mammary glands. **a**, Representative 3D surface rendering images of EpCAM-stained E13.5, E14.5, E15.5 and E16.5 epithelial mammary rudiments, based on 3D confocal imaging. Mammary gland 2 is shown. Scale bar, 20 µm. **b, c**, Quantification of epithelial mammary gland volume **(b)** and number of branching tips **(c)**, n_E13.5_=15, n_E14.5_=24, n_E15.5_=41, n_E16.5_=36. **d**, Confocal optical sections of whole mount-stained mammary glands from E13.5, E14.5, E15.5 and E16.5 Fucci2a embryos stained with EpCAM. Scale bars, 20 µm (E13.5-E15.5) and 30 µm (E16.5). **e**, Quantification of the proportions of all epithelial cells in S/G2/M and G1/G0 phases. Altogether, 15 glands (in total 9228 cells) from three E13.5 embryos, 24 glands (in total 17599 cells) from five E14.5 embryos, 41 glands (in total 40431 cells) from eight E15.5 embryos, and 36 glands (in total 50574 cells) from seven E16.5 embryos were analyzed. Data are presented as mean ± SD. **f**, A schematic image illustrating how the distance of cells (center of the nucleus) was quantified with respect to the surface of mammary rudiments. **g**, Density plot showing the distribution of the distance of nuclei in S/G2/M and G1/G0 phase to the surface of the mammary rudiment. Density plot revealed that a cluster of cells was localized within the distance of 10 µm (dashed line), which was set as the threshold to define “basal” and “inner” (luminal) cells. **h**, Quantification of the proportion of epithelial cells in S/G2/M phase in basal and inner compartments in E13.5-E16.5 epithelial mammary rudiments. Sample sizes are as in **e**. Data are presented as mean ± SD. The statistical significance was assessed using unpaired two-tailed Student’s *t*-test with Bonferroni correction. ns, non-significant; *, *p* < 0.05, ***, *p* < 0.001; ****, *p* < 0.0001.

Next, we asked whether the mesenchyme is needed for initiation of branching. To this end, we utilized a mesenchyme-free 3D mammary organoid technique to culture micro-dissected intact mammary rudiments in a serum-free medium with growth supplements ^21^(Fig. 1f). In the 3D Matrigel matrix, E16.5 mammary epithelia generated large branching trees in just 3 days (Fig. 2g,f and Supplementary Fig.1), whereas epithelia from earlier stages (E13.5 to E15.5) consistently failed to branch even after eight days of culture. Some specimens enlarged in size, yet they failed to progress, except for occasional E15.5 epithelia that generated a few branches (Fig. 2g,i and Supplementary Fig. 1).

Besides confirming previous observations ^19^ our results reveal that mesenchymes from advanced developmental stages could not alter the pace of epithelial outgrowth, yet epithelial-mesenchymal interactions are indispensable for the mammary epithelium to acquire branching ability.

### Basal-cell biased proliferation is activated in mammary epithelium prior to initiation of branching

Next, we sought to determine which mammary epithelial properties are required for the onset of branching. Majority of mammary epithelial cells are quiescent at the placode and bud stages ^22–24^, and proliferation is thought to resume when branching begins at around E16 ^22^. Such coincidence suggests that activation of proliferation may closely cooperate with, or even drive the onset of branching. To gain more insight into the quiescent stage of the embryonic mammary primordium, we first quantified the volume of the mammary epithelium with the aid of 3D surface renderings of EpCAM-stained specimens (Fig. 2a). The volume of mammary rudiments steadily increased from E13.5 to E16.5 (Fig. 2b), whereas quantification of the branch (tip) number showed that active branching did not take place until E16.5 (Fig. 2c).

To analyze epithelial proliferation between E13.5 and E16.5, we investigated cell cycle dynamics using the Fucci2a mouse model, where cells in S/G2/M phase of the cell cycle express nuclear mVenus while cells in G1/G0 express nuclear mCherry ^25^. The ratios of mammary epithelial cells in S/G2/M and G1/G0 phases were quantified in 3D after whole-mount staining with EpCAM (Fig. 2d). In line with the previous report ^24^, only ∼20% of mammary epithelial cells were in S/G2/M phase at E13.5, with no apparent change at E14.5 (Fig. 2e). However, the proportion of S/G2/M cells significantly increased at E15.5 but plateaued and even slightly decreased at E16.5 when branching was evident (Fig. 2e).

Initiation of branching morphogenesis has also been proposed to be associated with the onset of basal and luminal lineage segregation at E15-E16 ^8^. A recent single-cell RNA sequencing (RNAseq) profiling study identified a cluster of proliferative cells in addition to the luminal and basal cell clusters in the E15.5 mammary gland ^26^. These findings prompted us to examine whether the proliferative cells display any bias in their distribution at E13.5-E16.5. Due to the absence of clear spatial segregation of basal and luminal lineage markers during these early developmental stages ^9^, we instead measured the distance of each nucleus to the surface of epithelial mammary rudiments in 3D to define their location (Fig. 2f). Distribution of all nuclei revealed a cluster of cells localizing within 10 µm distance from the epithelial surface (dashed line in Fig. 2g), corresponding well with the confocal images showing radially organized, basally-located elongated cells in the same position (Fig. 2d,f). Next, we stratified the epithelial cells to basal (nuclear distance less than or equal to 10 µm from the surface) and inner (“luminal”) (nuclear distance more than 10 µm) ones and quantified the ratios of S/G2/M and G1/G0 cells in each compartment (Fig. 2h). At E13.5 and E14.5, the proportion of S/G2/M cells was higher in the inner compartment, though the difference was statistically significant only at E14.5. However, concomitant with the overall increase in proliferation (Fig. 2e), there was a switch in the proportion of S/G2/M and G1/G0 cells at E15.5 and E16.5, basal cells being significantly more proliferative.

### Basal-cell biased proliferation is not sufficient to drive initiation of branching

The observation that basal cell-biased proliferation occurred prior to onset of branching suggests that it might be a prerequisite for branching to occur. Interestingly, activation of epithelial cell proliferation also seems to be associated with basal cell lineage specification. To further investigate the potential link between lineage segregation, proliferation, and initiation of branching, we took advantage of a mouse model that displays precocious onset of branching, the K14-*Eda* mouse overexpressing Eda under the keratin 14 (K14) promoter. Eda and its epithelially-expressed receptor Edar regulate growth and branching of the embryonic and pubertal mammary gland ^13, 14, 27, 28^. In K14- *Eda* embryos, mammary epithelial proliferation is increased, and branching is initiated already at E14.5 ^13^.

To more closely examine the cellular alterations induced by Eda, we quantified the size, branch tip number, and proliferation status in K14-*Eda* embryos and their wild type littermates at E13.5 and E14.5. Mammary buds of K14-*Eda* embryos were significantly larger already at E13.5 (Fig. 3a,b), and at E14.5, the volume was comparable to those of E16.5 wild type embryos (compare Fig. 3b to Fig. 2b, all mice in C57Bl/6 background). As reported ^13^, branching was evident in K14-*Eda* embryos already at E14.5 (Fig. 3c).

**Fig.3.**
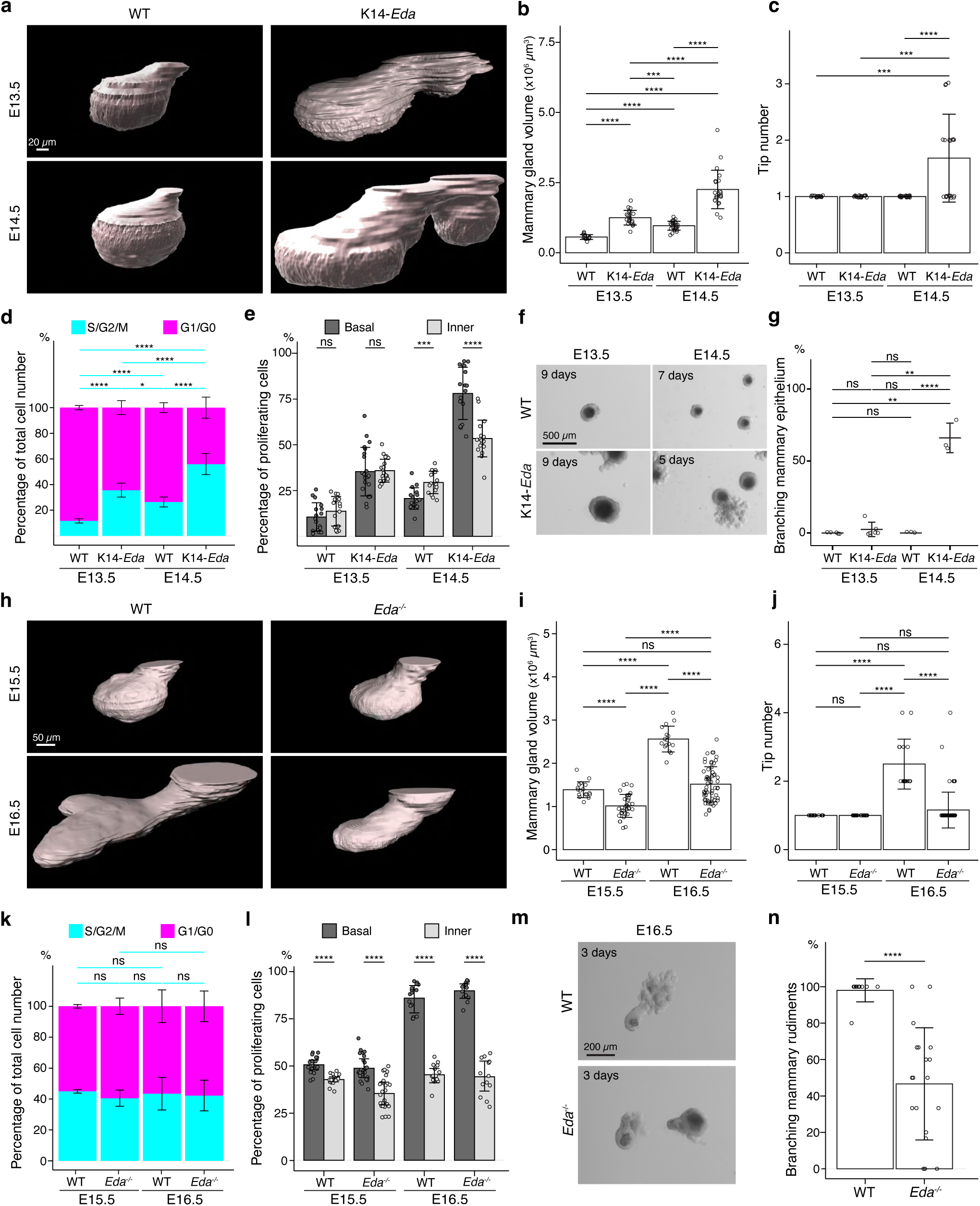
Basal-cell biased proliferation precedes, but is not sufficient to drive onset of branching. **a**, Representative 3D surface rendering images of EpCAM-stained mammary glands of K14-*Eda* embryos and their wild type (WT) littermates at E13.5 and E14.5. Mammary gland 2 is shown. Ectopic mammary rudiments (asterisk) common in K14-*Eda* embryos were excluded from the analysis. Scale bar, 20 µm. **b, c**, Quantification of mammary gland volume **(b)** and branching tip number **(c)** at E13.5 (n_WT_= 17, n_K14-Eda_= 21) and at E14.5 (n_WT_= 16 and n_K14-Eda_= 18). **d, e**, Quantification of the proportions of mammary epithelial cells in S/G2/M and G1/G0 phases in the entire epithelium (**d**) and the proportions of mammary epithelial cells in S/G2/M phase in basal and inner compartments (**e**) in WT or K14-*Eda* embryos at E13.5 (n_WT_= 17 glands and in total 7714 cells from 3 embryos, n_K14-Eda_= 21 glands and in total 15561 cells from 4 embryos) and E14.5 (n_WT_= 16 glands and in total 10221 cells from 4 embryos, n_K14-Eda_= 18 glands and in total 10520 cells from 5 embryos). **f**, Representative images showing the growth of E13.5 and E14.5 K14- *Eda* and wild type littermate epithelial mammary rudiments in 3D Matrigel culture. Note branching in E14.5 K14-*Eda* mammary rudiments. Scale bar, 500 µm. **g**, Quantification of branching mammary rudiments in 3D culture. Data are presented as percentage of branching mammary rudiments (mean ± SD) from a total of 4 (E13.5 WT), 6 (E13.5 K14-*Eda*), 3 (E14.5 WT) and 3 (E14.5 K14-*Eda*) independent experiments (each with minimum 5 rudiments in culture). **h**, Representative 3D surface rendering images of EpCAM-stained E15.5 and E16.5 epithelial mammary rudiments of *Eda^-/-^* and wild type embryos. Mammary gland 2 is shown. Scale bar, 50 µm. **i, j**, Quantification of epithelial mammary gland volume **(i)** and number of branching tips **(j),** at E15.5 (n_WT_ = 17 and n_Eda-/-_ = 27) and at E16.5 (n_WT_ = 16 and n_Eda-/-_ = 13). **k, l**, Quantification of the proportions of mammary epithelial cells in S/G2/M or G1/G0 phases (**k**) and the proportions of mammary epithelial cells in S/G2/M phase in basal and inner compartments (**l**) in WT or *Eda^-/-^* embryos at E15.5 (n_WT_ = 17 glands and in total 14054 cells from 3 embryos, n_Eda-/-_ = 27 glands and in total 21986 cells from 5 embryos) and E16.5 (n_WT_ = 16 glands and in total 40036 cells from 3 embryos, n_Eda-/-_ = 13 glands and in total 22009 cells from 3 embryos). **m**, Representative images showing E15.5 and E16.5 *Eda^-/-^* and wild type epithelial mammary rudiments in 3D culture after 3 days. Scale bar, 200 µm. **n**, Quantification of branching mammary rudiments in 3D culture. Data are presented as percentage of branching mammary rudiments from a total of 6 WT and 18 *Eda^-/-^* E16.5 embryos (each with 3-6 rudiments in culture). Data are presented as mean ± SD. The statistical significance was assessed using unpaired two-tailed Student’s *t*-test with Bonferroni correction. ns, non-significant; *, *p* < 0.05, **, *p* < 0.01, ***, *p* < 0.001; ****, *p* < 0.0001.

To specifically assess the link between lineage segregation and onset of branching, we analyzed expression of the well-established basal and luminal markers, keratin 14 (K14) and keratin 8 (K8) respectively, in K14-*Eda* and littermate control mammary glands (Supplementary Fig. 2a) between E13.5 and postnatal day 5 (P5). At E13.5, there was no obvious difference between the genotypes. Intriguingly, at E14.5 and E15.5, K14-*Eda* mammary rudiments displayed a notable downregulation of K8 not only in basal cells, but also in inner cells (Supplementary Fig. 2a). However, K8 expression resumed at E16.5, and at P1, and P5, no difference between the genotypes was observed (Supplementary Fig. 2a). Accordingly, gene set enrichment analysis of a microarray dataset of E13.5 *Eda^-/-^* mammary buds treated with recombinant Eda protein for 4 hours ^29^ revealed a positive enrichment of ‘LIM_Mammary_Stem_Cell_Up’ gene signature ^30^, known to represent the basal cell signature (Supplementary Fig. 2b). These data suggest that lineage segregation is unlikely to be a prerequisite for branching to commence, as branching was observed even though luminal lineage was transiently suppressed in K14-*Eda* embryos. However, we cannot exclude the possibility that consolidation of basal fate in basally located cells (by downregulation of luminal fate) plays a role.

Next, we focused on investigating the link between proliferation and onset of branching. Analysis of Fucci2a reporter expression in K14-*Eda* embryos at E13.5 and E14.5 revealed that the portion of S/G2/M cells was significantly higher in K14-*Eda* mice at both stages compared with wild type littermates (Fig. 3d and Supplementary Fig. 3a). In addition, the basal cell-biased proliferation was evident already at E14.5 (but not yet at E13.5) in K14-*Eda* embryos (Fig. 3e), similar to wild type mice at E15.5/E16.5 (Fig. 2h). Since E14.5 K14-*Eda* mammary glands had similar characteristics to E16.5 wild type in terms of volume, elevated overall proliferation, and basal cell-biased proliferation, we next tested their ability to grow and branch in the mesenchyme-free 3D Matrigel culture. E14.5, but not E13.5, K14-*Eda* epithelia were able to branch, whereas epithelia isolated from wild type littermates expectedly failed to generate outgrowths (Fig. 3f, g). We also analyzed Fucci2a reporter expression in *Eda*^-/-^ mice ^31^ at E15.5 and E16.5. As we previously reported ^13^, loss of *Eda* led to smaller glands and the onset of branching was delayed with most mammary glands being unbranched at E16.5 (Fig. 3h-j). However, the overall proliferation (Fig. 3k), as well as the relative portion of S/G2/M cells in basal and inner cells (Fig. 3l and Supplementary Fig. 3b) were similar between *Eda*^-/-^ and wild type controls at both stages. To evaluate the branching ability, we again performed mesenchyme-free 3D culture. While nearly all E16.5 control epithelia gave rise to branched outgrowths, as expected, about half of *Eda*^-/-^ epithelia failed to do so (Fig. 3m,n).

Collectively, these data indicate that initiation of branching succeeds activation of proliferation but is not its direct consequence. Additionally, analysis of K14-*Eda* mammary rudiments suggests that exit from quiescence and branching ability can occur independently of lineage specification.

### Salivary gland mesenchyme is rich in growth-promoting cues, but does not alter the mode of branch point formation of the mammary epithelium

Next, we shifted our focus on the regulation of the branching pattern, which is thought to be determined by mesenchymal cues ^19, 20^. To assess the influence of the mesenchyme, we performed heterotypic and heterochronic epithelial-mesenchymal recombination experiments between fluorescently labeled mammary and salivary gland tissues. Mammary epithelia and mesenchymes were isolated either at the quiescent bud stage (E13.5), or right after the bud had sprouted (E16.5); in addition to the primary mesenchyme, also mammary fat pad precursor tissue was micro-dissected from E16.5 embryos. Salivary gland tissues were isolated at E13.5, when the first branching events are evident and tissue separation is effortless. Homotypic recombinations were used as controls.

As previously reported ^19^, E16.5 mammary ductal trees were far denser when cultured with salivary gland mesenchyme, and grew and branched at a faster rate than with any of the mammary mesenchymes tested (Fig. 4a, top row). Of E13.5 mammary epithelia, majority (13 out of 18) did not survive in the salivary gland mesenchyme, and in the remaining ones, only traces of epithelial cells could be detected after 6 days of culture (Fig. 4a, middle row). However, E13.5 mammary epithelia branched readily in combination with all mammary mesenchymes (Fig. 4a, middle row), although their success rate was generally lower than that of E16.5 epithelia, as also previously reported ^19^. In addition, we assessed the impact of mammary mesenchyme on salivary gland epithelium. Although the salivary gland epithelium usually survived, further growth and branching were minimal when cultured with any of the mammary mesenchymes, in stark contrast with homotypic control recombinants (Fig. 4a, bottom row).

**Fig. 4.**
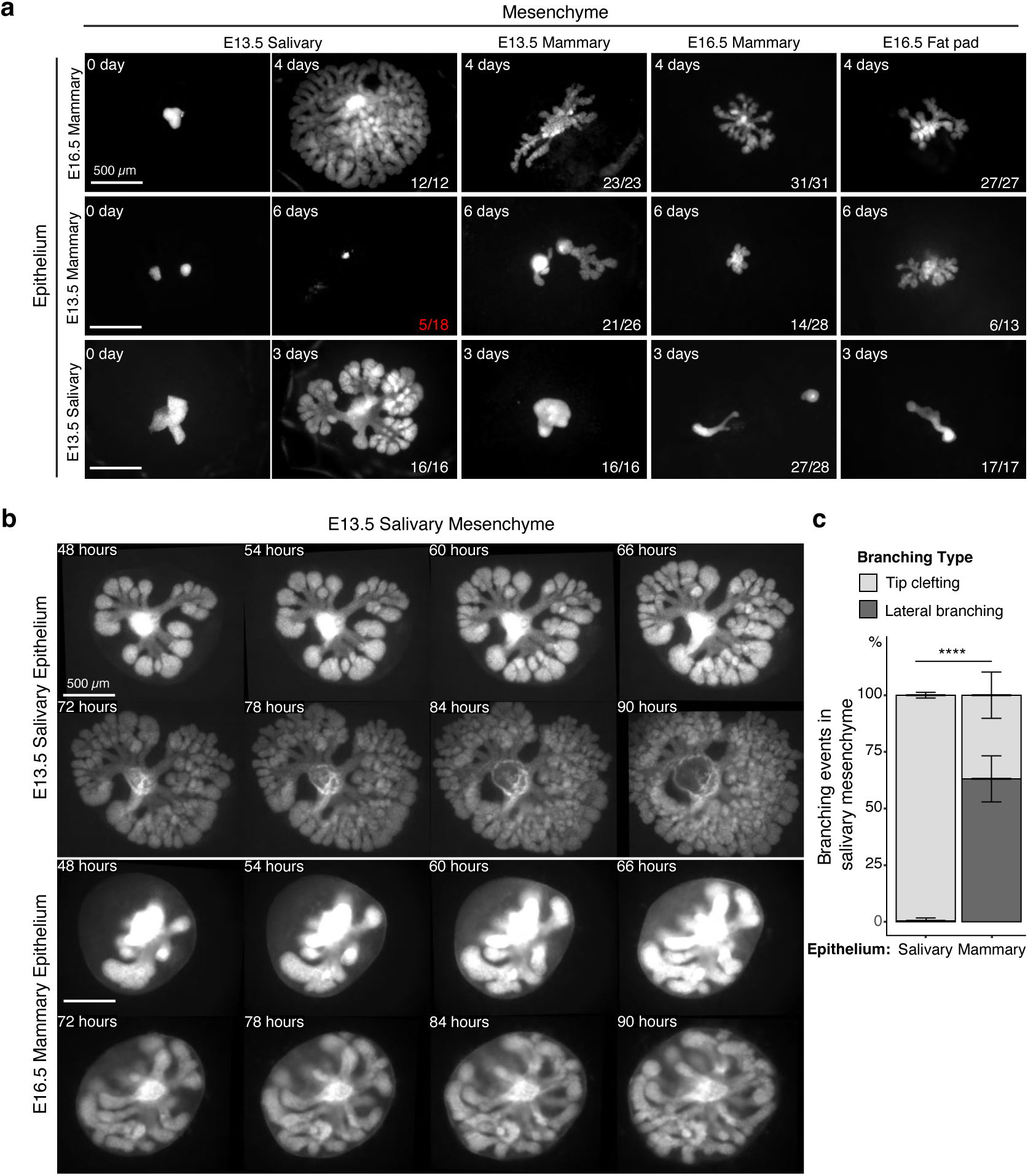
Mammary mesenchyme is indispensable for the branching ability of the mammary gland. Recombination experiments between micro-dissected mammary and salivary gland tissues using fluorescently labeled epithelia (see also Fig.1). **a**, Representative images showing growth of the indicated epithelia recombined with distinct mesenchymes. Images were taken 0-6 days after culture as indicated in each figure. n in the lower right corner indicates growing recombinants out of those that survived, except for E13.5 mammary epithelium recombined with E13.5 salivary gland mesenchyme where it shows the number of survived recombinants/total recombinants (in red). In these recombinants, the epithelia never branched. Data were pooled from 3-4 independent experiments. Scale bars, 500 µm. **b**, Captions of time-lapse live imaging series of explants consisting of E13.5 salivary epithelium or E16.5 mammary epithelium recombined with E13.5 salivary mesenchyme. Images were captured every 2 hours starting 48h after recombination. The full video is provided as Supplementary Video 1. Scale bar, 500 µm. **c**, Quantification of the branching events (lateral branching and tip clefting) from time-lapse videos. A pooled data from three independent experiments: in total of 239 branching events from 9 control recombinants (salivary epithelium + salivary mesenchyme) and 159 branching events from 8 explants consisting of mammary epithelium and salivary gland mesenchyme were analyzed. Data are represented as mean ± SD and the statistical significance was assessed with unpaired two-tailed Student’s *t*-test. *p* values: ****, *p* < 0.0001.

In principle, new branches can be generated by two different mechanisms: tip clefting/bifurcation or lateral (side) branching ^1, 3^. In the embryonic mammary gland, both events are common ^7^ while the salivary gland branches by tip clefting only ^32^. Recent advances in imaging technologies have enabled time-lapse analysis of branching events in detail prompting us to perform live imaging of salivary and mammary epithelia recombined *ex vivo* with salivary gland mesenchyme (Fig. 4b, Supplementary Video 1). Images were captured at 2h intervals, and branching events were traced and quantified from the time-lapse videos. Nearly all salivary gland branching events occurred by tip clefting (Fig. 4c), as expected. Surprisingly, over 60% of mammary branching events were generated by lateral branching, similar to normal embryonic mammary gland branching ^7^. We conclude that although salivary gland mesenchyme boosts growth of the mammary epithelium, the mode of branching is an intrinsic property of the mammary epithelium that is not altered by the growth-promoting salivary gland mesenchyme environment.

### Transcriptomic profiling of mammary and salivary gland mesenchymes identifies potential growth regulators

To identify the mesenchymal cues governing the differential growth characteristics of mammary and salivary gland epithelia, we performed transcriptomic profiling of five distinct tissues: E13.5 mammary mesenchyme surrounding the quiescent bud (E13.5 MM), E16.5 mammary mesenchyme surrounding the mammary sprout (E16.5 MM), E16.5 Fat pad precursor tissue (E16.5 FP), and E13.5 salivary gland mesenchyme (E13.5 SM) (Fig. 5a). E13.5 non-mammary ventral skin mesenchyme (E13.5 VM) was also included to allow identification of mammary-specific transcriptomes. Five biological replicates for each tissue were sequenced.

**Fig.5.**
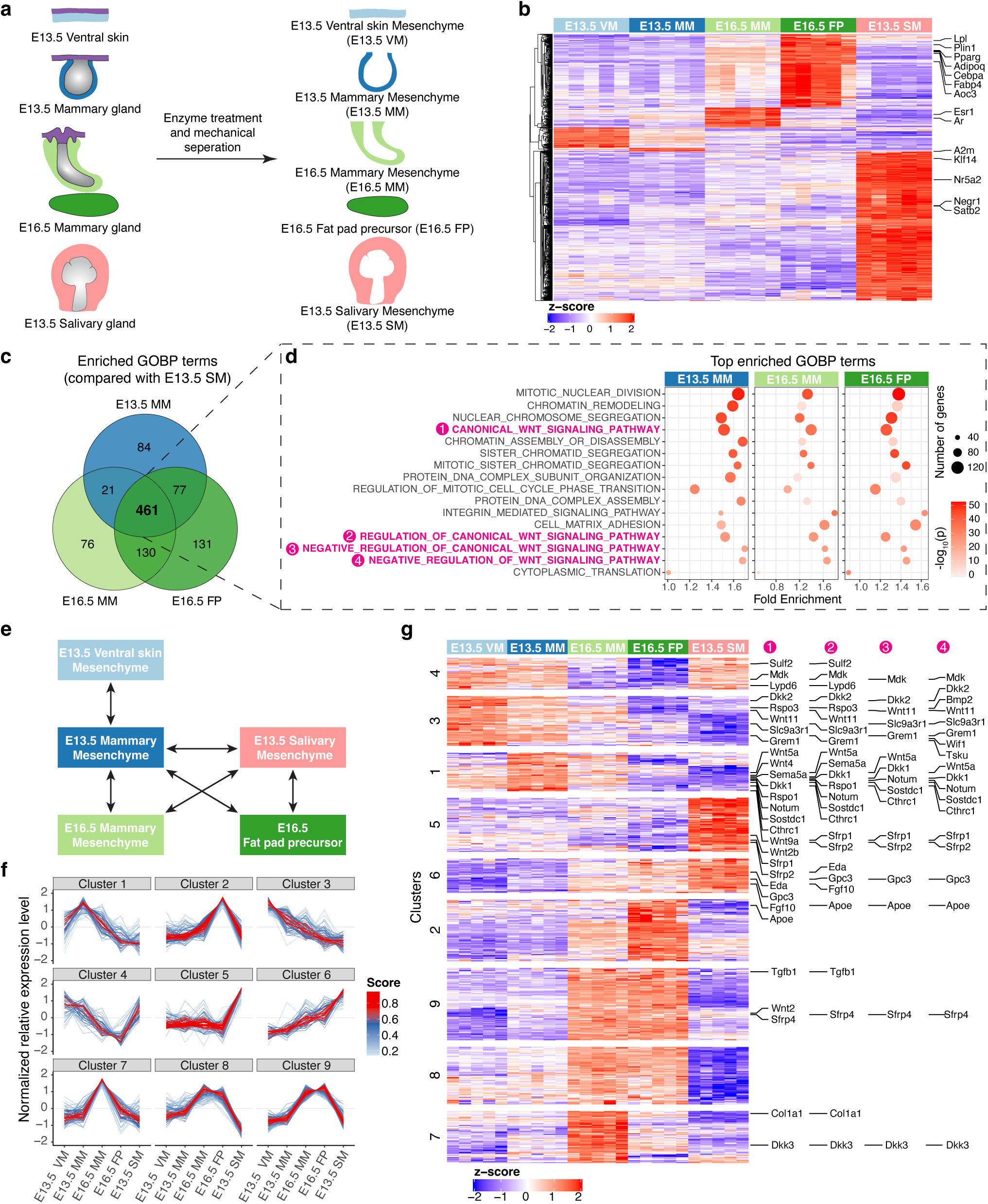
Transcriptomic analysis identifying mesenchymal signals potentially regulating epithelial growth. **a**, A scheme illustrating the tissues isolated for RNAseq analysis. **b**, Heatmap showing the expression of the identified marker genes (with a threshold of average of normalized expression value in each group ≥ 100, fold change ≥ 2 and adjusted p-value < 0.05) in different mesenchymes using the z-score of log2-transformed normalized expression value (also see Supplementary Table 1). **c**, Venn diagram showing 461 enriched Gene Ontology Biological Process (GOBP) terms shared among E13.5 mammary mesenchyme (MM), E16.5 MM and E16.5 fat pad (FP) when compared to E13.5 salivary gland mesenchymes (SM) separately. **d**, Top 10 (among the 461 shared terms) of the most significantly enriched GOBP terms in each comparison resulted in 16 distinct terms in total. Four out of 16 terms were related to Wnt signaling pathway (in magenta). **e**, A scheme illustrating the pair-wise comparisons used to identify the genes with the potential to regulate epithelial growth. Altogether 644 genes encoding extracellular matrix proteins and ligands with average of normalized expression value in each group ≥ 200, fold change ≥ 1.5 and adjusted p-value < 0.05 were identified. **f**, mFuzzy cluster analysis of the genes identified in **e**. **g**, Heatmap showing the expression of genes identified in **e** using the z-score of log2-transformed normalized expression value. The clusters were defined by mFuzzy shown in **f**. The genes within the Wnt related GOBP terms identified in **d** are indicated accordingly in the right.

The principal component analysis revealed that each group of samples were distinct from each other, although the E13.5 MM and E13.5 VM group quite close together (Supplementary Fig. 4a). To investigate the differences between the samples and assess the quality of the data, we performed pairwise comparisons and identified 51, 10, 54, 195, and 393 signature genes preferentially expressed in only one of the five sample sets (Fig. 5b and Supplementary Table 1). Among them, *Esr1* and *Ar* encoding estrogen and androgen receptors, respectively, were markers of E16.5 MM, while E16.5 FP was rich with adipogenesis markers such as *Aoc3*, *Adipoq*, *Cebpa*, *Fabp4, Lpl, Plin1 and Pparg* ^33^. E13.5 SM-enriched genes *Nr5a2*, *Negr1*, *Klf14* and *Satb2* have been identified as salivary mesenchyme markers by Sekiguchi et al. using single-cell RNA sequencing ^34^. These data indicate that our RNAseq data represent well the transcriptomes of the designated tissues.

To understand the functional disparity between salivary and mammary mesenchymes in promoting epithelial growth and branching, we performed a Gene Ontology (GO) enrichment analysis for differentially expressed genes (DEGs) in Biological Processes (BP) (Fig. 5c,d). In total, 461 GOBP terms were shared among E13.5 MM, E16.5 MM and E16.5 FP when compared to E13.5 SM. Among the 461 shared GOBP terms, the top 10 most significantly enriched terms in each pairwise comparison resulted only into 16 unique GOBP terms. Strikingly, of these, four were Wnt pathway related terms: canonical Wnt signaling pathway, regulation of canonical Wnt signaling pathway, negative regulation of Wnt signaling pathway, and negative regulation of canonical Wnt signaling pathway (Fig. 5d).

To identify genes with the potential to regulate epithelial cell behaviors, we focused on DEGs encoding extracellular (secreted or membrane-bound) molecules (signaling molecules, signaling pathway inhibitors, extracellular matrix components) in biologically relevant pairwise comparisons (Fig. 5e). Exclusion of lowly expressed genes led to the identification of 644 candidate genes (Supplementary Table 2). mFuzz cluster analysis ^35^ suggested that those genes could be further classified into 9 clusters based on their expression pattern across all the samples (Fig. 5f and Supplementary Table 2). Examination of the Wnt pathway related genes (as identified by GOBP enrichment analysis shown in Fig. 5d) in these clusters revealed that altogether 12 out of 19 negative regulators of Wnt pathway were markers of clusters 1 and 3, including *Dkk2*, *Bmp2*, *Wnt11*, *Slc9a3r1*, *Grem1*, *Wif1*, *Tsku*, *Wnt5a*, *Dkk1*, *Notum*, *Sostdc1* and *Cthrc1* (Fig. 5g). Clusters 1 and 3 were characterized by genes displaying lower expression in E16.5 MM than E13.5 MM, and the lowest level in E13.5 SM (Fig. 5f). Our tissue recombination experiments (Fig. 1b) suggest that such an expression pattern might represent potential growth suppressors. In other words, low expression of these negative regulators in salivary gland mesenchyme might enhance epithelial growth and branching, and in turn their higher expression in mammary mesenchyme might inhibit growth.

Clusters 2, 7, 8 and 9 were defined by genes such as *Hgf*, *Ltbp1, Tnc, and Postn,* with highest expression levels in one or more mammary-derived mesenchymes, highlighting them as best candidates to possess mammary-specific functions, e.g. in regulation of sprouting or epithelial cell differentiation. On the other hand, the clusters 5 and 6 genes, such as *Adam10*, *Adamts1*, *Bmp1*, *Bmp7*, *Fgf10*, *Igf1*, *Igf2* and *Eda,* have highest expression levels in E13.5 SM, indicating a potential role as drivers of epithelial growth. This fits well with the known roles of Eda and Fgf10 in salivary and mammary gland development ^7, 13, 36–38^.

### Wnt-activated mesenchyme promotes growth of the mammary epithelium

The transcriptomic analysis suggests that one significant difference between salivary and mammary mesenchymes is the Wnt pathway. Gene set variation analysis (GSVA) confirmed that the Wnt signaling signature was higher in E13.5 SM compared to all mammary mesenchymes (Supplementary Fig. 4b), which is consistent with the high expression of Wnt inhibitors in the mammary mesenchyme. Moreover, we have previously shown that suppression of mesenchymal Wnt activity in developing salivary glands compromises growth of the salivary gland ^36^. Together, these findings promoted us to ask whether low levels of mesenchymal Wnt activity could limit the growth of the mammary epithelium. To answer this question experimentally, we aimed to activate Wnt signaling by stabilizing β-catenin in the mesenchyme by crossing *Dermo1-Cre*^+/-^ mice with those harboring exon3 –floxed ß-catenin (*ß-catenin^flox3/flox3^* mouse) ^39^. However, this led to embryonic lethality already at E12.5, in line with previous reports ^40^. Therefore, we chose the tissue recombination approach where E13.5 wild type mammary buds were recombined with E13.5 mammary mesenchyme dissected either from control (*ß-catenin^wt/wt^*) or *ß-catenin^flox3/wt^* embryos, followed by adeno-associated virus (AAV8) – mediated gene transduction as a means to deliver Cre recombinase ^41^ (Fig. 6a). As a result, Wnt signaling was activated in the mesenchymal cells only. Quantification of tissue recombinants transduced with AAV8-Cre revealed that wild type mammary epithelia cultured on mammary mesenchyme from *ß-catenin^flox3/wt^* embryos had significantly more ductal tips than those cultured on control mammary mesenchyme (Fig 6b,c). These data indicate that low level of mesenchymal Wnt signaling activity limits growth and branching of the mammary epithelium.

**Fig.6.**
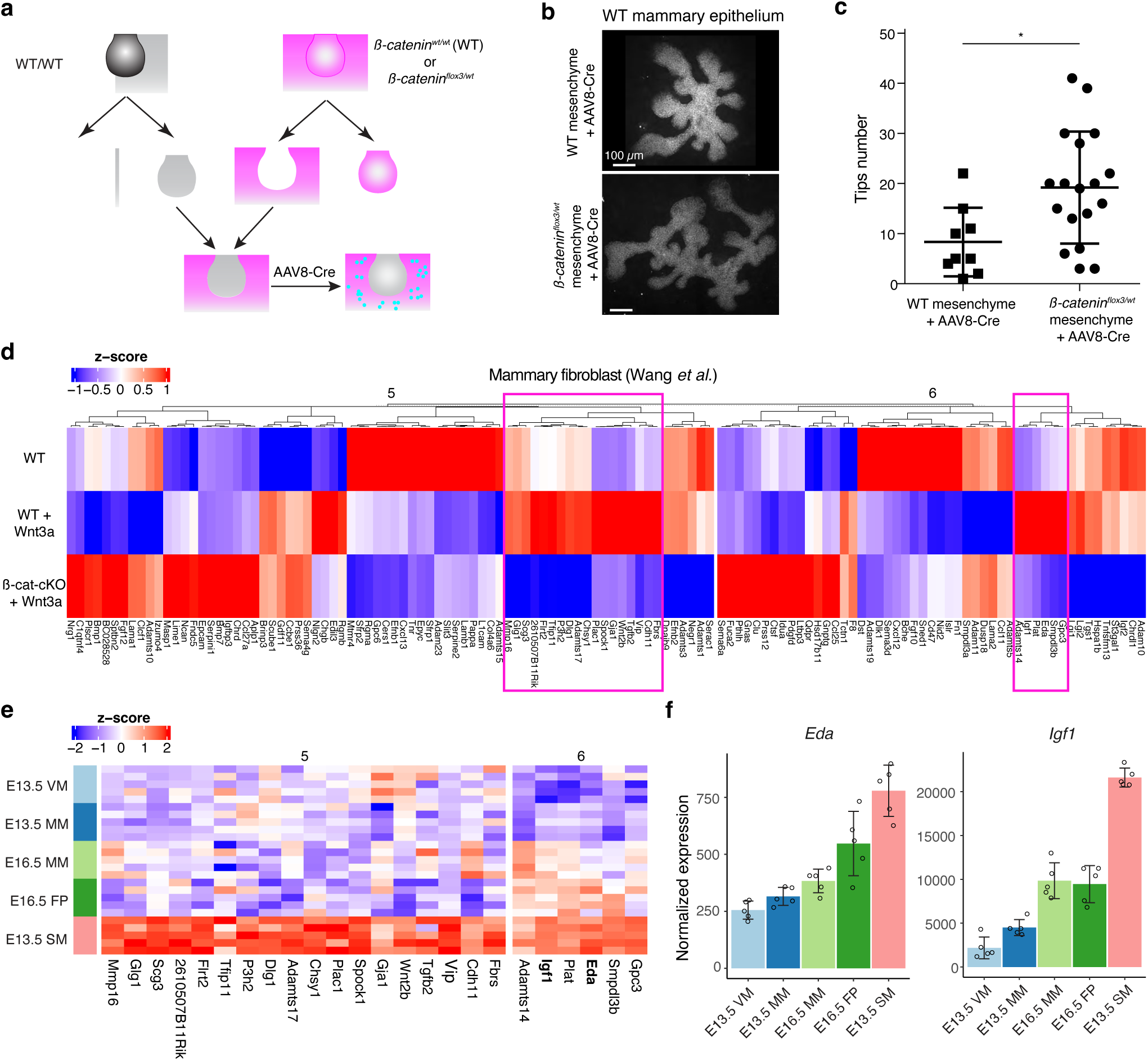
Wnt-activated mesenchyme promotes growth of the mammary epithelium. **a**, A scheme illustrating the experimental design for mesenchymal activation of Wnt/ß-catenin signaling activity. **b**, Representative images showing EpCAM stained wild type mammary epithelia after 6 days culture in wild type or *ß-catenin^flox3/wt^* mesenchyme infected with AAV8-Cre virus during the first 48 hours. **c**, Quantification of the number of branching tips of wild type mammary epithelia recombined with wild type or *ß-catenin^flox3/wt^* mesenchyme after 6 days of culture. Data are presented as mean ± SD (n = 9 and 18 for WT and *ß-catenin^flox3/wt^* mesenchyme, respectively) and represented from three independent experiments. Statistical significance was assessed using unpaired two-tailed Student’s *t*-test. *, *p* < 0.05. **d**, Unsupervised cluster of heatmap showing the expression of cluster 5 and 6 genes identified by mFuzzy analysis (see Fig. 5f) in a published dataset ^42^ that compared gene expression levels in wild type and *β-catenin* deficient mammary fibroblasts cultured with or without Wnt3a protein. Data are shown as z-score of log2-transformed normalized expression values. Two subsets of potential mesenchymal Wnt target genes identified are marked (box in magenta). **e**, Heatmap showing the expression of the candidate genes from **d** in different mesenchymes of the RNAseq data. Data are shown as z-score of log2-transformed normalized expression values. **f**, Graphs representing mRNA expression of *Eda* and *Igf1* as measured by RNAseq. Data are presented as normalized expression values (mean ± SD). Each dot represents one biological replicate.

Next, we asked which paracrine factors could regulate epithelial growth downstream of mesenchymal Wnt signaling. First, we explored a publicly available RNA-seq dataset ^42^ (Fig. 6d) which compared gene expression levels in wild type and β-catenin deficient mammary fibroblasts cultured with or without Wnt3a protein, and narrowed our analysis on cluster 5 and 6 genes identified in the mFuzz analysis (Fig. 5f and Supplementary Table 2). These genes displayed opposite expression patterns to genes in clusters 1 and 3, and hence were expected to positively regulate epithelial growth (Fig. 5f,g). The analysis revealed that the expression of most of the cluster 5 and 6 genes was altered in mammary fibroblasts upon manipulation of Wnt signaling activity (Fig. 6d). Focusing on genes upregulated by Wnt3a in wild type, but not in β-catenin deficient fibroblast led to the identification of 18 and 5 candidate genes in clusters 5 and 6, respectively, *Eda* and *Igf1* being amongst them (Fig. 6d-f). We have previously identified *Eda* as a gene downstream of Wnt pathway in the salivary gland mesenchyme ^36^, validating our analysis pipeline.

### IGF-1R is required for embryonic mammary gland development and branching morphogenesis

IGF-1 is well known for its role in growth control and, similar to other tissues, it functions as an important local mediator of the growth hormone in pubertal mammary glands ^43–45^. However, the role of the IGF-1 pathway in embryonic mammary gland development has not been explored, apart from one study reporting the smaller size of the E14 mammary bud in IGF-1R deficient embryos ^46^. Analyses of the known secreted components of the IGF pathway revealed that many of them were differentially expressed between salivary and mammary gland mesenchymes (Supplementary Fig. 5), the most striking being *Igf1* and pregnancy-associated plasma protein-A (*Pappa*), a zinc metalloproteinase that promotes IGF-1 signaling through cleavage of the inhibitory Igf-binding proteins (IGFBPs) ^47^. *Pappa* was also identified as a cluster 5 gene in the mFuzz analysis (Supplementary Table 2). To functionally test the effect of IGF-1 on mammary gland growth, we performed *ex vivo* culture of E16.5 mammary glands and treated the explants for 3 days with moderate levels of recombinant IGF-1 or vehicle (Fig. 7a). Quantification of branch tip number showed that IGF-1 significantly increased growth of the mammary epithelium (Fig. 7b).

**Fig. 7.**
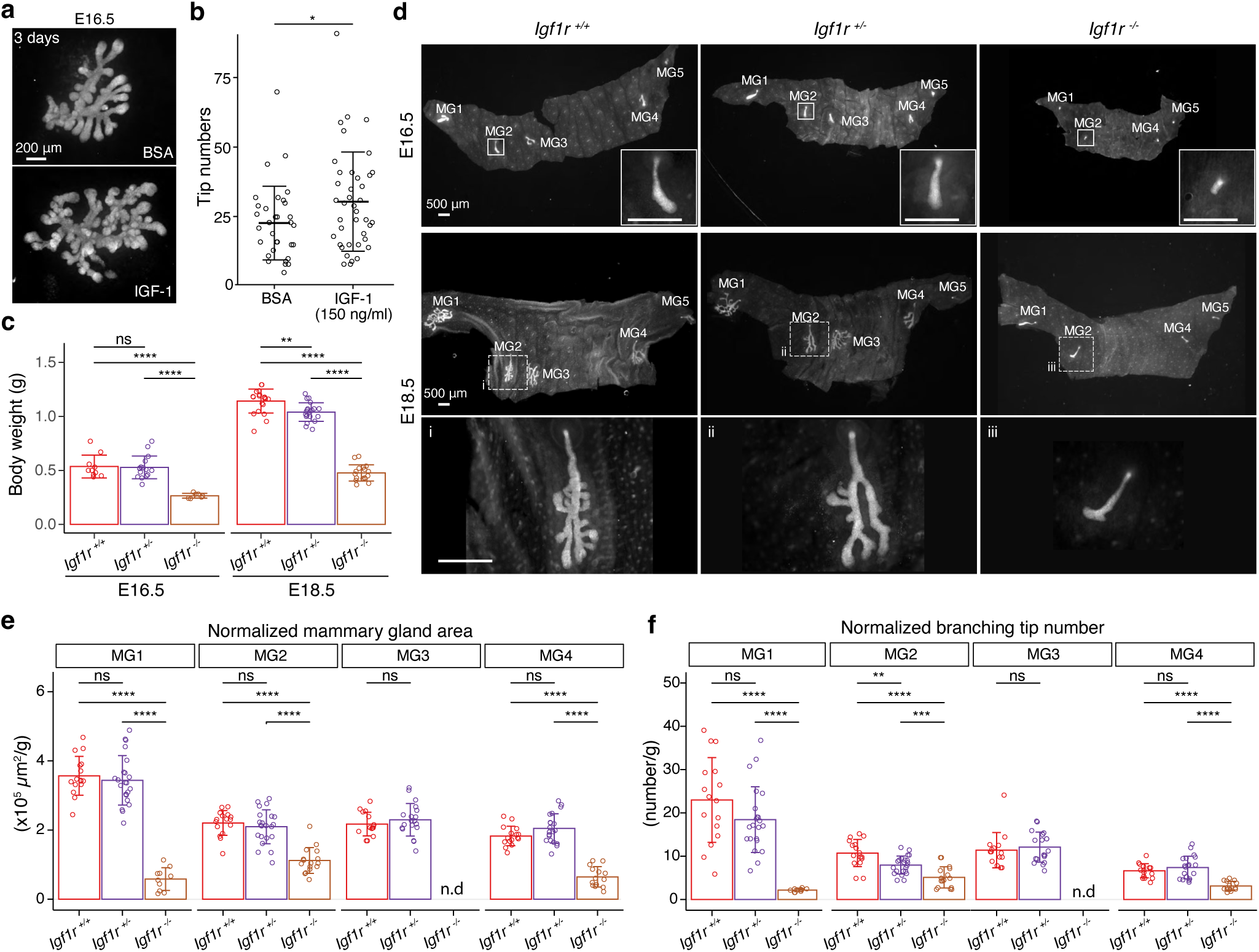
IGF-1R is required for embryonic mammary gland development and branching morphogenesis. **a**, Representative images of E16.5 K14-Cre::mTmG mammary glands cultured *ex vivo* for three days in the presence of 150 ng/ml recombinant IGF-1 or vehicle (BSA). Scale bar, 200 µm. **b**, Quantification of the number of branching tips in vehicle (n=33) and IGF-1 treated (n=40) mammary gland explants. Data are pooled from 5 independent experiments and presented as mean ± SD. Statistical significance was assessed using unpaired two-tailed Student’s *t*-test. *, *p* < 0.05. **c**, Body weight of *Igf1r^+/+^*, *Igf1r^+/-^* and *Igf1r^-/-^*embryos at E16.5 (n*_Igf1r+/+_*=10, n*_Igf1r+/-_*=16; n*_Igf1r-/-_*=7), and E18.5 (n*_Igf1r+/+_*=20, n*_Igf1r+/-_*=20; n*_Igf1r-/-_*=17). Data are presented as mean ± SD. Statistical significances were calculated with unpaired two-tailed Student’s *t*-test with Bonferroni correction. ns, non-significant; **, *p* < 0.01; ****, *p* < 0.0001. **d**, Representative images of EpCAM-stained ventral skin including mammary glands (MG) 1-5 from *Igf1r^+/+^*, *Igf1r^+/-^* and *Igf1r^-/-^* female embryos at E16.5, and E18.5. Note absence of MG3 in *Igf1r^-/-^* embryos. Magnifications show mammary gland 2. Scale bars, 500 µm. **e, f**, Quantification of mammary gland area **(e)** and number of branch tips **(f)** normalized to body weight in *Igf1r^+/+^*, *Igf1r^+/-^* and *Igf1r^-/-^* embryos at E18.5. MG5 was often lost during dissection and therefore was not included in the analysis. n.d, not detected. Data are presented as mean ± SD and the statistical significances were assessed using unpaired two-tailed Student’s *t*-test with Bonferroni correction. ns, non-significant; *, *p* < 0.05, **, *p* < 0.01, ***, *p* < 0.001; ****, *p* < 0.0001.

To assess the function of IGF-1 *in vivo*, we examined mammary gland development in embryos deficient for *Igf1r*, the obligate cognate receptor of Igf1 ^48, 49^. As previously reported ^50^, *Igf1r^-/-^* embryos were significantly smaller compared with wild type littermates (*Igf1r^+/+^)* (Fig. 7c). At E16.5, the anterior glands of littermate control embryos had sprouted. Small outgrowths were also observed in *Igf1r^-/-^* embryos, with the exception of mammary gland 3 that was consistently absent (Fig. 7d). At E18.5, growth and branching was severely compromised in the *Igf1r^-/-^* embryos, verified by quantification of the epithelial area of the mammary gland and the ductal tip number of mammary glands 1-4 at E18.5 (Supplementary Fig. 6a,b). To avoid biases caused by the conspicuously smaller size of the *Igf1r^-/-^* embryos ^50, 51^, we normalized the data to the body weight (Fig. 7e,f). The normalized values revealed that the mammary gland area and tip numbers were significantly reduced in *Igf1r^-/-^* embryos compared to controls. There was no significant difference between *Igf1r^+/-^* and *Igfr1^+/+^* embryos, except that the number of tips in mammary gland 2 was reduced in *Igf1r^+/-^* embryos (Fig. 7f). Analysis of E13.5 embryos revealed that mammary rudiment 3 was absent in *Igf1r^-/-^* embryos already early on (Supplementary Fig. 6c,d). We also examined the developing salivary glands at E13.5, E16.5 and E18.5. In stark contrast to the mammary gland, the salivary glands of E16.5 and E18.5 *Igf1r^-/-^* embryos were highly branched although smaller (Supplementary Fig. 6e), paralleling the overall growth defect of the mutant embryos (Fig. 7c). Collectively, these data show that embryonic mammary gland development is exceptionally sensitive to loss of IGF-1/IGF-1R signaling, as shown by the complete absence of mammary bud 3 and the specific growth and branching impairment during late embryogenesis.

## Discussion

In this study, we explored the fundamental principles of epithelial-mesenchymal tissue interactions guiding embryonic mammary gland development. Our findings reveal that while both the timing and type of branching events are intrinsic properties of the mammary epithelium, mammary-specific mesenchymal signals are crucial for the acquisition of the branching capacity. Importantly, we demonstrate that salivary gland mesenchyme could only promote the growth of mammary epithelium without changing the branching regime. Transcriptomic profiling and experimental evidence indicate that mesenchymal Wnt signaling and Igf1 downstream of it are critical regulators promoting mammary gland growth and contribute to the differences in growth-promoting capacity of the mammary and salivary mesenchymes. Other pathways are also involved, as several signaling molecules known to regulate growth, such as *Eda* and *Fgf10* ^7, 52, 53^, were differentially expressed between salivary and mammary gland mesenchymes.

Three important events occur before initiation of mammary gland branching: exit from quiescence, lineage segregation and initial outgrowth. Our data suggest that these three phenomena are likely coordinated, in part through Eda signaling. Interestingly, basal cells are more proliferative initially, unlike during later embryogenesis when branching is ongoing ^54^. Our observation that basal cell-biased proliferation is activated prior to branching seems to support the previous hypothesis that proliferation and lineage segregation may be prerequisites for branching ^8, 10^. However, basal cells are more proliferative also in K14-*Eda* mammary glands where establishment of luminal fate is temporarily disrupted when branching begins. This finding argues that lineage segregation is independent of onset of branching. Moreover, analyses in both Eda loss- and gain-of-function mice imply that even though cell proliferation precedes, it alone is not sufficient to induce branching. This is in line with our recent study showing that inhibition of cell proliferation does not prevent branch point generation or branch elongation *per se*, though new cells are evidently needed as building blocks for further ductal growth ^54^. Instead, cell motility is critical for branch point formation in the mammary gland ^54^, as well as in other branching organs ^55–57^.

Epithelial-mesenchymal tissue recombination experiments performed mainly in the 50’s to 70’s using different branched organs, including the lung, kidney, and salivary gland, have disclosed the dominant role of the mesenchyme in branch patterning ^17, 58–64^, a conclusion confirmed also by detailed branch pattern analyses of heterotypic kidney and lung tissue ^18^. Similarly, recombination experiments between mammary epithelium and salivary gland mesenchyme ^19, 20^ laid the foundation for our current understanding on the instructive role of the mesenchyme in mammary gland branching morphogenesis. However, at the time, time-lapse imaging was not feasible precluding a comprehensive investigation of the dynamic branching process. Advances in imaging may explain our contrasting result. That is, our data clearly demonstrate that although the density and growth rate of the mammary ductal tree were greatly enhanced by the salivary gland mesenchyme, the type of branch point formation was not. This observation suggests that mammary epithelium itself carries the instructions dictating the mode of branching involving both lateral branching and tip bifurcations. This conclusion is further supported by our recent study showing that isolated E16.5 mammary epithelia retain bimodal branching also in the mesenchyme-free 3D organoid culture ^54^. Evidently, further studies are required to elucidate which properties of the mammary epithelium enable its bimodal branching behavior.

In contrast to the mode of branching, growth rate and density of the mammary ductal tree was grossly altered by the salivary gland mesenchyme implying an important role for paracrine factors in these processes. This, together with the failure of the salivary epithelium to grow in mammary gland mesenchyme indicate that the mammary gland mesenchyme is either poor in growth-promoting cues and/or rich in growth-inhibitory cues. Our transcriptomic profiling suggest that it may be both. Our RNA-seq data indicated that low level of mesenchymal Wnt activity may restrict mammary gland growth. Mesenchymal Wnt activity is critical for the early specification of the mammary mesenchyme ^65^, but its function beyond the bud stage is largely unknown. Here, our experimental data revealed that growth and branching of the mammary gland was enhanced by mesenchymal activation of Wnt/β-catenin signaling activity. Previous studies have shown that an excess of Wnt ligands promotes growth of the embryonic mammary epithelium but the primary target tissue was unknown ^13, 66^. Our results suggest that this could be (in part) an indirect effect, due to augmented mesenchymal Wnt signaling activity.

The IGF-1/IGF-1R signaling pathway has a critical role in the coordinate regulation of body growth downstream of the pituitary growth hormone ^49, 67^. In its absence, the size of the organs is also proportionally reduced ^49, 68^. Here we show that the embryonic mammary gland is particularly sensitive to *Igf1r* deficiency, mammary gland 3 failing to develop at all. These data suggest that the role of IGF-1R during mammary gland development, particularly in the branching morphogenesis, extends beyond its general growth promoting function during embryonic development. The reason for this is currently unknown but one possibility is that the availability of active IGFs in mammary gland mesenchyme is limited to begin with, due to low expression of *Pappa*. Normally, the IGFs exist in the form of binary complexes with IGFBPs, and PAPPA degrades IGFBPs, increasing the bioavailable fraction of IGFs thereby promoting activation of IGF-1R ^49^.

In conclusion, our findings provide valuable insights into the growth control of the mammary gland and the transcriptomic profiling of different mesenchymes as a novel resource for investigating the mesenchymal contribution in organ development. Intriguingly, we found that heterochronic mammary mesenchyme did not advance/delay the timing of epithelial outgrowth and branching, indicating that mechanisms intrinsic to the mammary epithelium govern these processes. Yet, mammary-specific mesenchyme was indispensable for branching to occur, suggesting that mammary mesenchyme may provide permissive cues that allow the mammary bud to exit quiescence and become competent to respond to mitogenic cues. Parathyroid hormone like hormone (Pthlh, also known as Pthrp) signaling may play a critical role here: deletion of the mesenchymally expressed receptor *Pthr1* or the epithelially expressed ligand halts mammary gland development at E15.5-E16.5, prior to onset of branching ^69^. However, the downstream targets of Pthr1 are incompletely understood, but both Wnt and bone morphogenetic protein (Bmp) pathways are involved ^65, 70^. In addition, the transcriptomic and epigenetic changes taking place in the mammary epithelium between the quiescent bud stage and growth competent sprout are currently unknown. Uncovering how mammary epithelial cells acquire their remarkable growth potential and identification of the underlying mesenchymal cues are fascinating avenues for future research with implications to our understanding of basic breast biology, as well as breast cancer.

## Methods

### Mice

To obtain mice constitutively expressing the Fucci2a cell cycle reporters (R26R-Fucci2a-del/del), the conditional R26R-Fucci2a-floxed/floxed (Fucci2a^floxed/floxed^) mice ^25^ were first bred with PGK-Cre mice ^71^ ubiquitously expressing Cre. The obtained PGK-Cre;Fucci2a^del/floxed^ offspring were used to generate Fucci2a (R26R-Fucci2a-del/del) mice without the PGK-Cre transgene. Heterozygous R26R-Fucci2a-del/wt embryos were used for the quantitative analysis. The dual fluorescent mGFP;mTmG (R26-mGFP;mTmG) mice were generated by breeding mTmG (R26R-mTmG/mTmG) mice (ICR background; the Jackson Laboratory Stock no. 007576) with mGFP (R26R-mGFP/wt) mice (mixed background). The mGFP allele was generated by breeding mTmG mice with PGK-Cre mice ^71^ to remove the sequence containing the mTdtomato coding region and STOP cassette surrounded by loxP sites leading to ubiquitous expression of mGFP. The obtained PGK-Cre;mGFP mouse was bred with wild type C57Bl/6 mouse to remove the PGK-Cre transgene. For embryonic tissue recombination experiments, male mGFP;mTmG mice were mated with wild type NMRI females. K14-*Eda* and *Eda^-/-^* mice were maintained as described previously ^13^. The K14-*Eda*;Fucci2a embryos were obtained by crossing K14- *Eda* males with Fucci2a-del/del females. As the *Eda* gene is localized in the X-chromosome, to obtain the Fucci2a;*Eda^-/-^* and Fucci2a;*Eda^+/+^* embryos, the Fucci2a mice were first bred with *Eda^-/y^* male or *Eda^-/-^* female to obtain Fucci2a;*Eda^+/y^* and Fucci2a;*Eda^-/y^* males, and Fucci2a;*Eda^+/-^* females. For the analysis, the Fucci2a;*Eda^-/-^* embryos were obtained by breeding Fucci2a;*Eda^-/y^* males with Fucci2a;*Eda^+/-^* females and Fucci2a;*Eda^+/+^* embryos were obtained by breeding Fucci2a;*Eda^+/y^* males with Fucci2a;*Eda^+/-^* females. The *ß-catenin^flox3/flox^*^3^ mice ^39^ were maintained in C57Bl/6 background as described previously ^72^. *ß-catenin^flox3/flox3^* or *ß-catenin^wt/wt^* (wild type C57Bl/6) male mice were bred with C57Bl/6 wild type females to obtain the *ß-catenin^flox3/wt^* or *ß-catenin^wt/wt^* embryos for the AAV virus transduction experiments*. Igf1r^+/-^* mice were maintained in 129S2/SvPasCrl background as described previously ^51^. The littermates obtained from breeding of *Igf1r^+/-^* male and *Igf1r^+/-^* female mice were used for analysis.

All mice were kept in 12 hours light-dark cycles with food and water given ad libitum. The appearance of the vaginal plug was considered as embryonic day 0.5, and the age of the embryos was further verified based on the limb and craniofacial morphology and other external criteria ^73^. For embryos older than E13.5, only female embryos were used for experiments and analysis. The gender was determined by the morphology of the gonad as described previously ^21^ and further confirmed by detecting the Y chromosomal *Sry* gene using PCR ^74^.

### *Ex vivo* embryonic tissue culture and tissue recombination

*Ex vivo* culture of embryonic mammary glands was performed as described earlier^21^. Briefly, the abdominal-thoracic skin containing mammary glands 1-3 was dissected from E13.5 to E16.5 embryos. The tissues were treated for 30-60 min with 2.5 U/ml of Dispase II (4942078001; Sigma Aldrich) in PBS at +4⁰C in the shaker and then 3-4 min with a pancreatin-trypsin (2.5 mg/ml pancreatin [P3292; Sigma Aldrich] and 22.5 mg/ml trypsin dissolved in Thyrode’s solution pH 7.4) at room temperature. The tissues were incubated in culture media (10% FBS in 1:1 DMEM/F12 supplemented with 100 μg/ml ascorbic acid, 10 U/ml penicillin and 10 mg/ml streptomycin) on ice for a minimum of 30 min before further processing. The skin epithelium was removed with 26-gauge needles leaving the mesenchymal tissue with the mammary buds.

For typical mammary gland culture, the tissues were collected on small pieces of Nuclepore polycarbonate filter with 0.1 µm pore size (WHA110605, Whatman) and further cultured on the air-liquid interface on filters with the support of metal grids in a 3.5 cm plastic Petri dish with culture medium. The explants were cultured in a humidified incubator at 37°C with an atmosphere of 5% CO_2_ and the culture medium was replaced every other day.

To test the role of Igf1 in branching morphogenesis, recombinant mouse IGF1 protein (791-MG, R&D systems) at the final concentration of 150 ng/ml was added to the culture medium 3 hours after the onset of the culture. The same volume of 10% BSA was used as a vehicle control. The fresh culture medium with IGF1 or BSA was replaced after two days, and the explants were cultured for three days in total.

For tissue recombination experiments, embryos expressing mGFP or mTmG were identified with a fluorescent stereomicroscope and processed separately. The E13.5 submandibular glands (hereafter salivary gland) were dissected and processed similarly as described above for the mammary gland. After enzyme treatment and incubation on ice, the tissues were further dissected under a stereomicroscope to separate the intact mammary or salivary gland epithelium and their mesenchyme. The mesenchymes without any epithelium were collected with the filter and maintained in the culture incubator until further use. For salivary mesenchyme, mesenchymes from 3-4 salivary glands were pooled into one piece of filter to increase the amount of mesenchyme in each sample. After epithelial-mesenchymal separation of all samples, salivary epithelium or mammary buds 1-3 were gently washed by pipetting through a 1000 μl tip several times to remove the remaining mesenchymal tissues and then transferred onto the mesenchyme expressing different fluorescent protein, as previously described ^41^. 1-2 mammary buds were transferred to each mesenchyme. The recombinants were cultured as described above.

To specifically activate the WNT/ß-catenin signaling in the mesenchyme, tissue recombination has been performed as described above, while the mesenchymes from E13.5 *ß-catenin^flox3/^*^wt^ or *ß-catenin^wt/wt^* embryos were recombined with mammary buds from *ß-catenin ^wt/wt^* embryos. 2 hours after culture, final concentration of 1.13×10^7^ vg/µl AAV8-Cre (purchased from AAV Gene Transfer and Cell Therapy Core Facility, Faculty of Medicine, University of Helsinki) were added into the culture medium. The fresh culture medium without virus was replaced every other day, and the explants were cultured 6-7 days in total.

### Time-lapse imaging for recombinants

To monitor the growth of the recombinants, the explants were imaged with Zeiss Lumar microscope equipped with Apolumar S 1.2x objective once per day. To assess the branching type of each event of the epithelium in salivary mesenchyme, multi-position, automated time-lapse imaging described previously ^21^ was used instead. Briefly, tissue recombination was performed as described above (Day 0). 1-2 days after the culture, explants with filter were transformed to 24 mm Transwell inserts with 0.4 µm polyester membrane (CLS3450, Costar) and cultured on 6-well plates allowing multi-position imaging ^7^. From day 1 or 2 to day 4 of culture, explants were imaged with 3i Marianas widefield microscope equipped with 10x/0.30 EC Plan-Neofluar Ph1 WD=5.2 M27 at 37 °C with 6% CO2. The medium was changed right before the imaging and thereafter, every other day. Images were acquired with an LED light source (CoolLED pE2 with 490 nm/550 nm) every 2 hours.

### Mesenchyme-free mammary rudiment culture and time-lapse imaging

E13.5 to E16.5 mammary rudiments were cultured in 3D Matrigel as previously described ^21^. Briefly, after separation of the mammary tissue with mesenchyme, the intact mammary rudiments 1-3 were dissected under stereomicroscope as described above. The mammary rudiments collected from littermate embryos of same genotype were pooled together, except for *Eda^-/-^* or *Eda^+/+^* . Pooled mammary rudiments 1-3 from each *Eda^-/-^* and *Eda^+/+^* embryo were cultured separately as it is not possible to obtain *Eda^-/-^* and *Eda^+/+^* genotypes from the same litter. Intact mammary rudiments were transferred onto the bottom of 12-well plates with 10 μl of culture media. The medium was then replaced with a 20-30 μl drop of growth-factor reduced Matrigel (356231; Corning) using a chilled pipette tip. The MBs were dispersed to avoid any potential contact with each other or the bottom of the plate. The mixture was then incubated in the 37°C culture incubator for 15-20 minutes until the matrix was solidified. The MBs were cultured in a humidified incubator at 37°C with an atmosphere of 5% CO_2_ in serum-free DMEM/F12 medium supplemented with 1X ITS Liquid Media Supplement (I3146, Sigma Aldrich] and 2.5 nM hFGF2 (CF0291; Sigma Aldrich), 10 U/ml penicillin and 10 000 μg/ml streptomycin. The culture medium was replaced every other day and the growth of the MBs was monitored once per day by imaging with Zeiss Lumar microscope.

### Whole-mount immunofluorescence staining and imaging

For whole-mount immunofluorescence staining, dissected ventral skin containing mammary glands, cultured explants, or mammary epithelia cultured in Matrigel were fixed in 4% PFA at 4°C overnight, washed three times in PBS and then three times in 1% PBST (1% TritonX-100 in PBS) at room temperature. Samples were blocked with blocking buffer containing 5% normal donkey serum, 0.5% BSA, and 10 μg/ml Hoechst 33342 (Molecular Probes/Invitrogen) in 1% PBST at 4°C overnight. The samples were then incubated with primary antibodies diluted in blocking buffer for 1-2 days at 4°C, washed three times with 0.3% PBST at room temperature before incubation with secondary antibodies diluted in 0.3% PBST with 0.5% BSA for 1-2 days at 4°C. After washing three times with 0.3% PBST and three times with PBS, samples were post-fixed with 4% PFA for 10 minutes at room temperature. Finally, samples were washed twice with PBS before immersing into the fructose-glycerol based clearing solution described by Dekker et al. ^75^ before imaging. For samples from older embryos, the blocking step was extended to 2 days followed by an extra microdissection procedure, where samples were dissected under fluorescence stereomicroscope to expose the mammary epithelium and remove surplus mesenchymal tissues. The samples were imaged with Leica TCS SP8 inverted laser scanning confocal microscope with HC PL APO 20x/0.75 IMM CORR CS2 object. The images were acquired with z-stack of 0.11 µm intervals.

For E13.5 *Igf1r* embryos, the staining was performed with the whole embryos before imaging. The samples of *Igf1r* embryos or IGF1-treated explants were imaged with Lumar stereomicroscope.

The following antibodies were used in this study: rat anti-mouse CD326 (EpCAM, 552370, BD Pharmingen, 1:500), rabbit anti-mouse Krt14 (RB-9020-P, Thermo Fisher Scientific, 1:500), rat anti-Krt8 (TROMA-1, DSHB, 1:500), Alexa Fluor 488-conjugated Donkey anti-Rat secondary antibody (A21208, Invitrogen, 1:500) and Alexa Fluor 647-conjugated Donkey anti-Rat secondary antibody (A48272, Invitrogen, 1:500).

### Image analysis

For mammary gland volume quantification, the border of mammary epithelium and mesenchyme was outlined manually based on EpCAM expression and bud morphology and the surface rendering and volume quantification were performed with Imaris 9.2 software (Bitplane). The mammary gland tip number was counted manually in 3D using Imaris. To further quantify the cell cycle dynamics of mammary epithelial cells, the mammary epithelium was masked using the rendered mammary gland surface in Imaris. Epithelial cells expressing nuclear mCherry (G1/G0) or nuclear mVenus (S/G2/M) were automatically detected using spot detection function with manual correction. The distance of each detected nucleus to the mammary epithelium surface was measured using the distance transformation function of Imaris. All the data were exported to be further analyzed using R version 4.2.1, a free software environment available at https://www.r-project.org/.

To quantify the mammary gland growth affected by the deficient of *Igf1r,* the epithelial area of the mammary glands and the number of ductal tips were acquired manually with ROI Manager within ImageJ (Fiji, version 1.53t) ^76^. -Time-lapse images were analyzed with ImageJ manually.

The plots were produced with R using packages tidyverse version 1.3.2 ^77^, ggplot2 version 3.4.0 ^78^, ggsignif version 0.6.4 ^79^, ggpubr version 0.4.0 ^80^ and RcolorBrewer version 1.1-3 ^81^.

### RNA sequencing and data analysis

To obtain the mesenchyme samples for RNA sequencing, salivary glands or flank skins with mammary rudiments 1-3 were dissected and followed by enzyme treatment as described above for *ex vivo* embryonic tissue culture. E13.5 salivary gland mesenchymes were obtained after removing the salivary gland epithelium. For E13.5 and E16.5 mammary gland mesenchymes, after removing the skin epithelium, the mammary epithelium and its surrounding mesenchyme were isolated together with small scissors followed by removal of the mammary epithelium using 26-gauge needles (303800, BD Microlance). The E16.5 fat pad precursor was microdissected from the explants after enzyme treatment. The E13.5 ventral skin mesenchymes further away from the mammary gland region were collected as E13.5 skin mesenchyme. The mesenchymes isolated from 2-3 embryos from the same litter were pooled together as one sample. Altogether, five biology replicates for each sample were collected from 3 different litters of C57Bl/6JOlaHsd mice. Samples were lysed immediately after collection and stored in TRI Reagent (T9424, Sigma) at -80°C. Total RNA was extracted using Direct-zol RNA Microprep kit (Zymo Research, Irvine, CA) with DNase treatment according to the manufacturer’s instructions. RNA quality was assessed with 2100 Bioanalyzer (Agilent, Santa Clara, CA) using Agilent RNA 6000 Pico Kit or Agilent RNA 6000 Nano Kit (Agilent, Santa Clara, CA). RNA concentration was determined using Qubit RNA HS Assay Kit (Q32855, Thermo Fisher) with Qubit 4 Fluorometer (Thermo Fisher). cDNA libraries were prepared with Ovation SoLo RNA-seq System (NuGen/Tecan Genomics) according to manufacturer’s instructions and sequenced with NextSeq 500 (Illumina, San Diego, CA) in the DNA Genomics and Sequencing core facility, Institute of Biotechnology, HiLIFE, University of Helsinki. 45-68 million reads per sample were obtained after 3 rounds of sequencing.

For RNAseq data analysis, all sequencing reads were processed for quality control, removal of low-quality reads, adaptor sequence and ribosomal RNA using fastqc version 0.11.8 ^82^, multiqc version 1.9 ^83^, Trimmomatic version 0.39 ^84^ and SortMeRNA version 2.1 ^85^ accordingly. The filtered reads were mapped to the reference genome (mm10) using Salmon version 0.99.0 ^86^ resulting in 36.6 to 53.4 million mapped reads per sample. The GSVA analysis was performed with R package GSVA version 1.44.5 ^87^. The conversion of murine gene Ensembl IDs to human Entrez IDs was performed with the biomaRt package version 2.46.3 ^88, 89^, using the reference mart https://dec2021.archive.ensembl.org. The significant differentially expressed signatures between different mesenchymes were assessed with lmFit and eBayes functions from R package limma version 3.52.4 ^90^, by comparing E13.5 MM, E16.5 MM or E16.5 FP with E13.5 SM, respectively. The signature database was downloaded from www.gsea-msigdb.org ^91^ on 12^th^ February 2023. The significantly enriched KEGG signaling pathways were pooled together for visualization. The data normalization and analysis of differentially expressed genes (DEGs) were performed using the R package DESeq2 version 3.15 ^92^. DEGs were defined with the thresholds of average count number > 50, adjusted p-value <0.05 and Log2(Fold Change) >=0.58 in each pairwise comparison.

Gene Ontology enrichment analysis was performed with the DEGs using R package pathfindR version 1.6.4 ^93^. Only the GOBP terms with lowest adjusted p value less than 0.01 were considered as significant. Among the commonly significantly altered GOBP terms, the top 10 GOBP terms with lowest adjusted p-value in each comparison and totally 16 GO terms were plotted. Gene Ontology database was downloaded from MSigDB ^91^ using R package msigdbr version 7.5.1 ^94^ on 9^th^ November 2022.

The DEGs with an average count number >100 and upregulated more than twice (Log2(Fold change)>=1) in each group of samples compared to all the other 4 groups of samples were identified as marker genes.

To detect the pattern of the gene expression among different mesenchymal tissues, DEGs encoding extracellular matrix protein or ligands in selected pairwise comparisons with an average count number >200 in each group were further analyzed using Mfuzz version 2.58.0 ^95^. The average of the normalized count number of each group was used as input. In addition, the groups were converted to pseudotime for the analysis. The fuzzifier m was determined with the default function and returned a value of 2.113207. The number of clusters was optimized empirically and set as 9 for the final analysis. The curated database including ECM, Ligand or Receptor genes was combined from the databases of R package SingleCellSignalR version 1.2.0 ^96^, CellTalkDB version 1.0 ^97^ and curated GO terms downloaded from https://baderlab.org/CellCellInteractions ^98^.

The plots were produced using R packages tidyverse version 1.3.2 ^77^, ggplot2 version 3.4.0 ^78^, circlize version 0.4.15 ^99^, RcolorBrewer version 1.1-3 ^81^, pathfindR version 1.6.4 ^93^, ComplexHeatmap version 2.12.1 ^100^, venn version 1.11 ^101^ and patchwork version 1.1.2 ^102^.

### Public RNAseq data analysis

The raw data from Wang et al. ^42^ (OEP001019) were downloaded from https://www.biosino.org/node/index. The sequence reads were processed similarly as described above. The log2 transformed normalized expression of selected genes were extracted to construct the heatmap shown in Fig. 6D.

### Statistical analysis

All data were analyzed by Prism 9 (GraphPad Software), or R packages ggsignif version 0.6.4 ^79^ and ggpubr version 0.4.0 ^80^. Statistical tests used are indicated in figure legends. p-values < 0.05 were considered significant. Throughout the figure legends: *p < 0.05, **p<0.01; ***p < 0.001, ****p < 0.0001.

### Ethics statement

All mouse experiments were approved by the Laboratory Animal Center at the University of Helsinki and the National Animal Experiment Board of Finland with the licenses number KEK19-019, KEK22-014 and ESAVI/2363/04.10.07/2017. Mice were euthanized with CO_2_ followed by cervical dislocation.

### Data availability

The raw and processed RNAseq data created in this study have been deposited in the GEO database under the access code GSEXXXXXX.

### Code availability

The code used for the analyses is open-source and available through the R packages described in the methods. All the customized scripts for producing the figures in this study are available upon request to the corresponding author.

## Acknowledgements

The authors wish to thank Dr. Jianpin Cheng, Dr. Alison Kuony and M.Sc. Aida Kaffash Hoshiar for the critical comments and suggestions on the manuscript, Ms. Raija Savolainen and Ms. Merja Mäkinen for excellent technical assistance, Dr. Maria Voutilainen, Dr. Satu-Marja Myllymäki and Dr. Ana-Marija Sulić for technical advice, past and present members of the Mikkola lab for insightful discussions. We also acknowledge Dr. Rishi Das Roy for the important discussion on RNAseq data analysis and CSC – IT Center for Science, Finland, for computational resources. AAVs were provided by AAV Gene Transfer and Cell Therapy Core Facility, Faculty of Medicine, University of Helsinki. Confocal and widefield microscope imaging and image analysis were performed at the Light Microscopy Unit, Institute of Biotechnology, supported by HiLIFE and Biocenter Finland. RNA sequencing was performed in the DNA Sequencing and Genomics Unit at the Institute of Biotechnology, HiLIFE, University of Helsinki. This work was carried out with the support of HiLIFE Laboratory Animal Centre Core Facility, University of Helsinki, Finland.

This work was supported by the Academy of Finland project grant (318287 to M.L.M.) and Center of Excellence Program (272280 and 307421 to M.L.M.), the Cancer Society of Finland (M.L.M.), the Jane and Aatos Erkko Foundation (M.L.M.), the Sigrid Jusélius Foundation (M.L.M.), the HiLIFE Fellow Program (M.L.M.), Oskar Öflunds Foundation (Q.L.), the Doctoral Programme in Integrative Life Science of the University of Helsinki (E.T.), the Doctoral Programme in Biomedicine (M.C.), the Finnish Cultural Foundation (J.S. and B.K.), and Ella and Georg Ehrnrooth Foundation (J.S.).

## Author Contributions

Conceptualization, Q.L., M.L.M.; Methodology, Q.L., R.L.; Software, Q.L.; Validation, Q.L.; Formal analysis, Q.L., J.S., B.K.; Investigation, Q.L., E.T., R.L., J.S., M.C.; Resources, M.H., J.J., M.L.M.; Data Curation, Q.L.; Writing – original draft preparation, Q.L., M.L.M; Writing – review and editing, Q.L., E.T., R.L., J.S., B.K., M.C., M.H., J.J., M.L.M; Visualization, Q.L.; Supervision, M.L.M.; Project administration, Q.L., M.L.M; Funding acquisition, M.L.M.

---

**Supplementary Fig.1.**
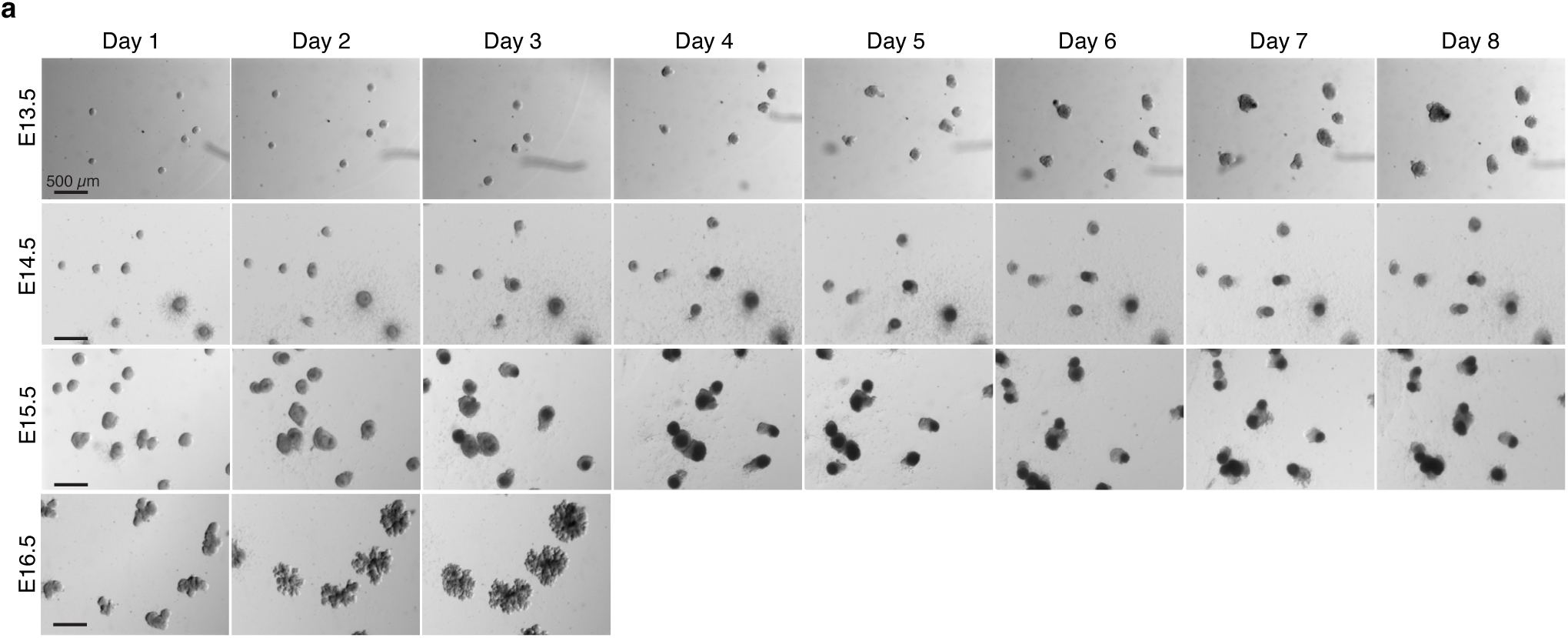
Epithelial mammary rudiments isolated from stages earlier than E16.5 are not able to branch in 3D culture. Representative images showing the growth of E13.5, E14.5, E15.5 and E16.5 epithelial mammary rudiments 1 in 3D Matrigel culture. Images were acquired once per day. Scale bar, 500 µm.

**Supplementary Fig. 2.**
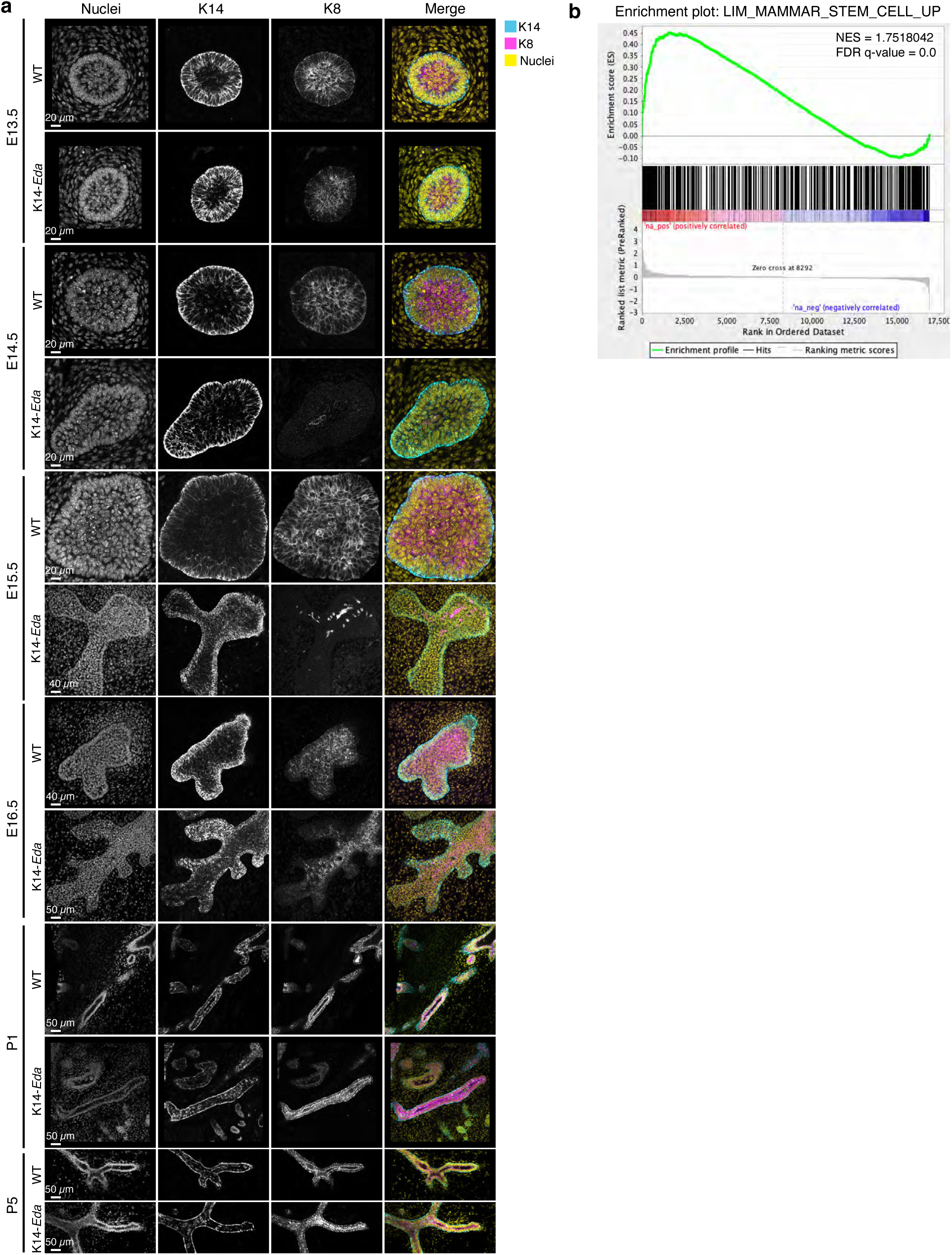
Overexpression of Eda transiently suppresses the luminal cell fate. **a**, Confocal optical sections of whole mount imaged E13.5, E14.5, E15.5, E16.5, postnatal day 1 (P1), and P5 wild type and K14-*Eda* mammary glands stained with the basal marker K14 (Cyan in merged images) and luminal marker K8 (Magenta in merged images). Nuclei were visualized with Hoechst. Mammary glands from at least 3 embryos from 2-3 litters were examined in each condition. Scale bars, 20-50 µm as indicated in the figures. **b**, Gene set enrichment analysis of Voutilainen data ^29^ of E13.5 *Eda^-/-^* mammary buds treated with recombinant Eda protein for 4 hours revealed a positive enrichment of ‘LIM_Mammary_Stem_Cell_Up’ gene signature ^30^.

**Supplementary Fig. 3.**
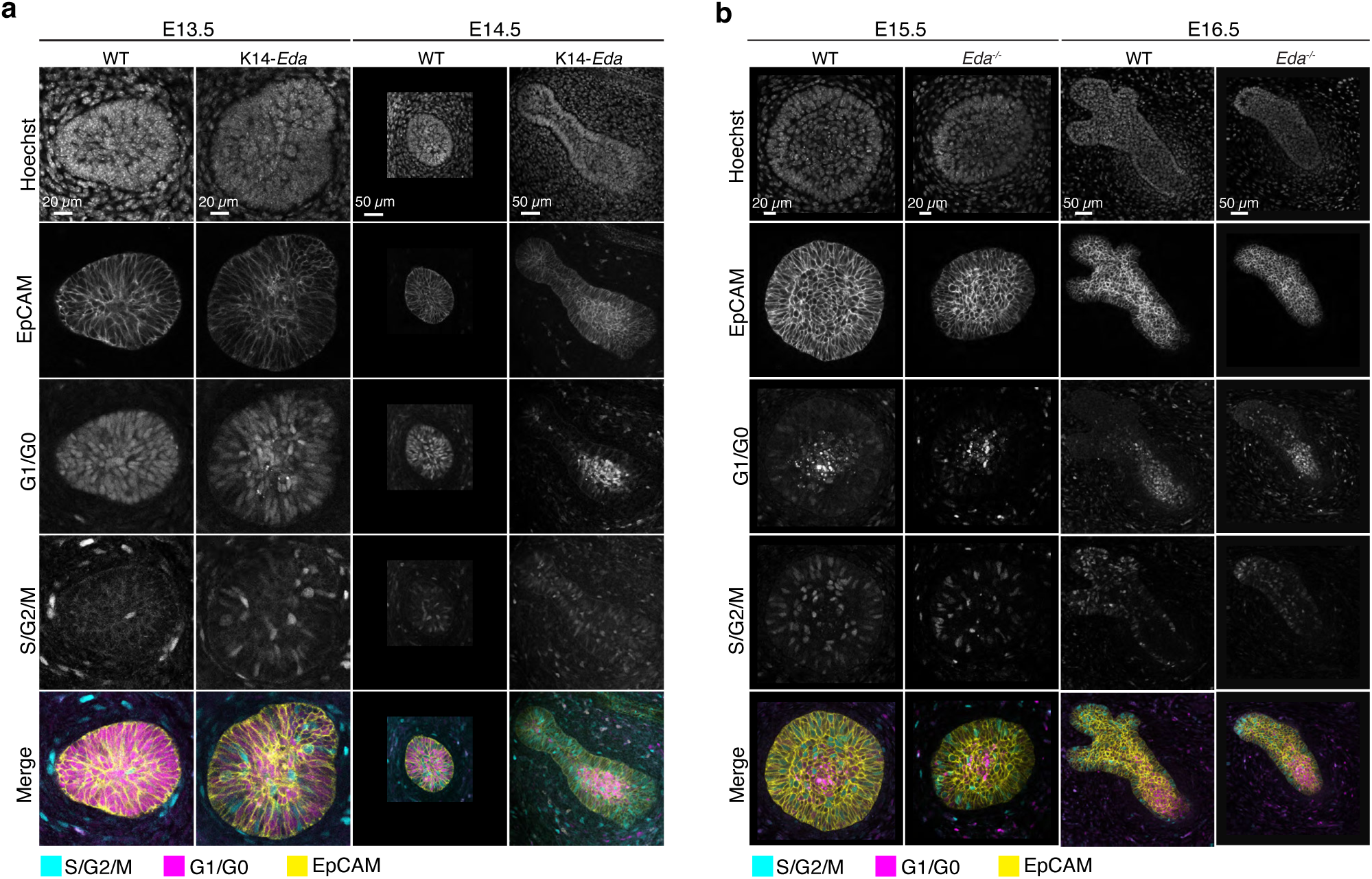
The proliferation dynamics of mammary epithelium in *Eda* gain-of-function and loss-of-function mouse models. **a**, Confocal optical sections of whole-mount mammary glands from E13.5 and E14.5 K14-*Eda* or WT littermate embryos expressing Fucci2a reporter stained with EpCAM. Scale bars, 20 µm (E13.5) and 50 µm (E14.5). **b**, Confocal optical sections of whole-mount mammary glands from E15.5 and E16.5 WT or *Eda^-/-^* Fucci2a embryos stained with EpCAM. Scale bars, 20 µm (E15.5) or 50 µm (E16.5).

**Supplementary Fig 4.**
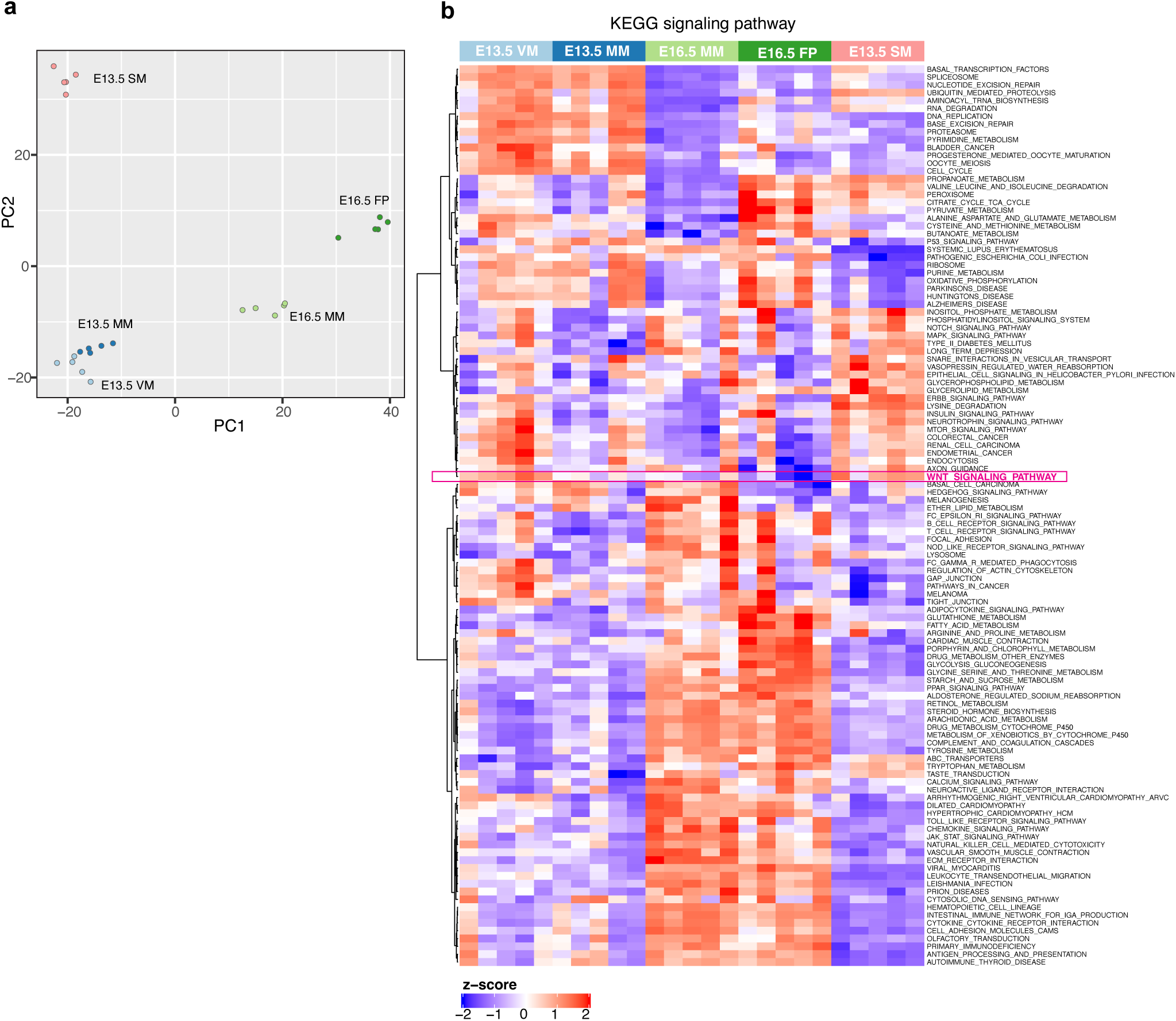
Transcriptomic profiling of different mesenchymes. **a**, Scatter plot shows the principal component analysis of E13.5 ventral skin mesenchyme (VM), E13.5 mammary mesenchyme (MM), E16.5 MM, E16.5 fat pad (FP), and E13.5 salivary gland mesenchyme (SM). **b**, Heatmap shows the significantly altered KEGG signaling pathways comparing E13.5 MM, E16.5 MM or E16.5 FP with E13.5 SM separately. WNT_SIGNALING_PATHWAY (marked with Magenta) is low in E16.5 MM and E16.5 FP compared to other tissues.

**Supplementary Fig. 5.**
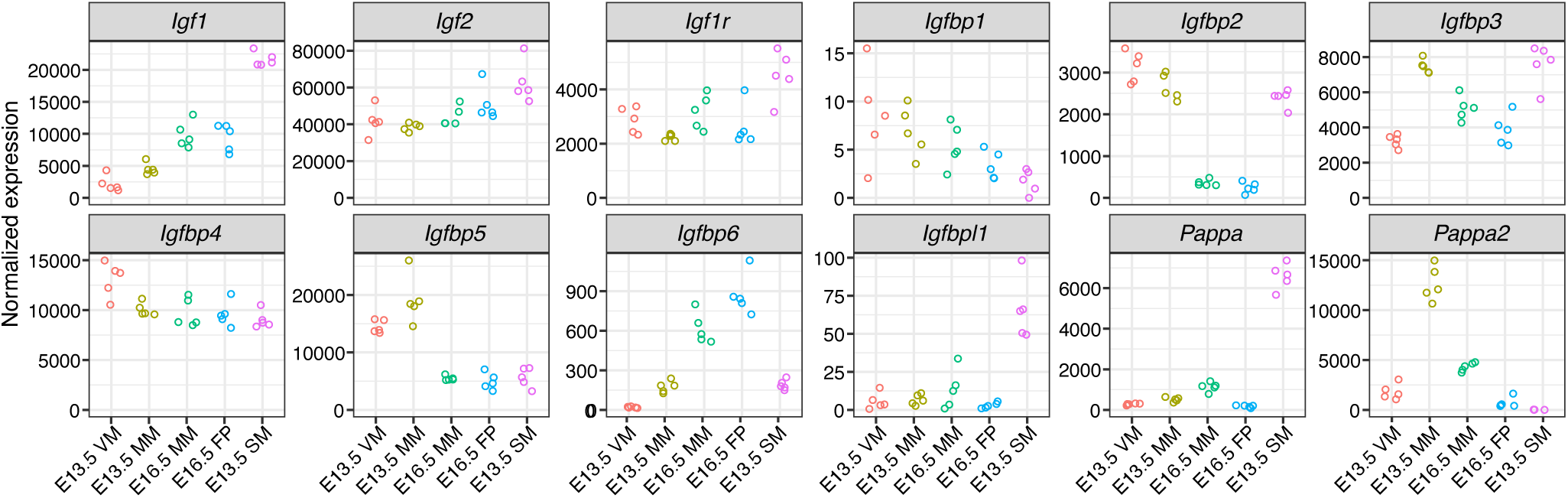
Expression of IGF pathway genes in the mesenchymal tissues. Graphs show mRNA expression of the indicated genes by RNAseq in E13.5 ventral, non-mammary skin mesenchyme (VM), E13.5 mammary mesenchyme (MM), E16.5 MM, E16.5 fat pad precursor (FP), and E13.5 salivary gland mesenchyme (SM). Each dot represents one biological replicate.

**Supplementary Fig. 6.**
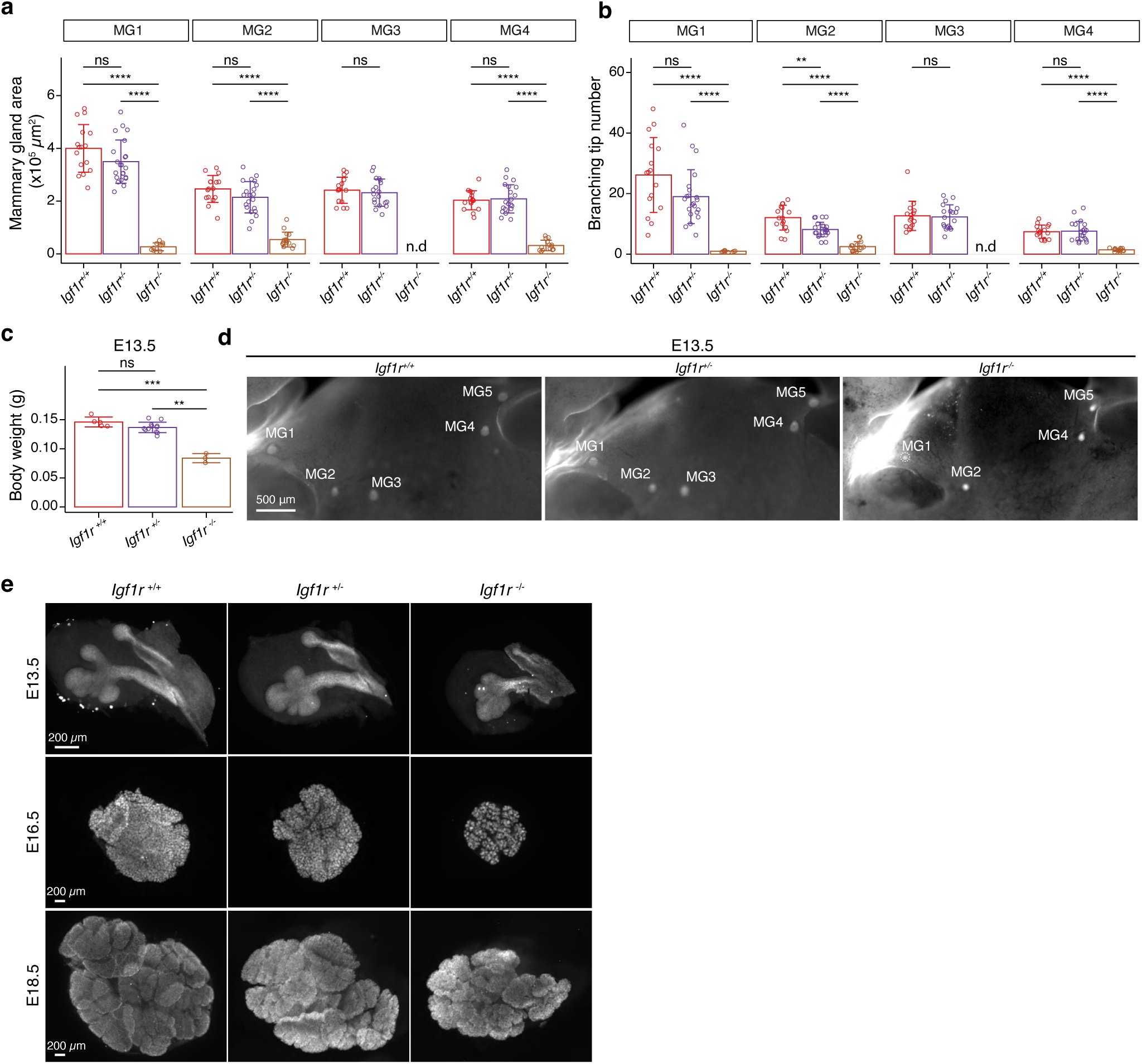
Impact of *Igf1r* deficiency on mammary gland and salivary gland growth and branching. **a, b,** Quantification of mammary gland area **(a)** and tip number **(b)** of E18.5 *Igf1r^+/+^*, *Igf1r^+/-^* and *Igf1r^-/-^* female embryos. Samples are the same as in Fig. 7e,f. Data are presented as mean ± SD. **c**, Body weight of *Igf1r^+/+^*, *Igf1r^+/-^*, and *Igf1r^-/-^* embryos at E13.5 (n*_Igf1r+/+_*=5, n*_Igf1r+/-_*=13; n*_Igf1r-/-_*=3). Data are presented as mean ± SD. **d**, Representative images of EpCAM-stained E13.5 embryos showing mammary glands (MG) 1-5 from *Igf1r^+/+^*, *Igf1r^+/-^* and *Igf1r^-/-^* embryos. Scale bar, 500 µm. **e,** Representative images of EpCAM-stained *Igf1r^+/+^*, *Igf1r^+/-^*, and *Igf1r^-/-^* salivary glands at E13.5 (n*_Igf1r+/+_*=6, n*_Igf1r+/-_*=8; n*_Igf1r-/-_*=20), E16.5 (n*_Igf1r+/+_*=15, n*_Igf1r+/-_*=29; n*_Igf1r-/-_*=5), and E18.5 (n*_Igf1r+/+_*=6, n*_Igf1r+/-_*=13; n*_Igf1r-/-_*=6). Scale bars, 200 µm. Statistical significances were assessed using unpaired two-tailed Student’s *t*-test with Bonferroni correction. ns, non-significant; *, *p* < 0.05, **, *p* < 0.01, ***, *p* < 0.001; ****, *p* < 0.0001.

**Supplementary Table 1.** The list of identified marker genes for each mesenchyme and their normalized expression value in each sample.

**Supplementary Table 2.** The results of mFuzzy analysis shown in Fig. 5f and the normalized expression value of each gene in each sample.

**Supplementary Video 1.** Time-lapse live imaging showing the growth of E13.5 salivary epithelium (left) and E16.5 mammary epithelium (right) in E13.5 salivary mesenchyme. Images were captured every 2 hours starting 48h after recombination. Scale bar, 500 µm.

